# Quantitative and amino acid sequence analysis of soluble HIV-1 Vpu and calmodulin interactions

**DOI:** 10.64898/2026.04.15.718738

**Authors:** Adeyemi Ogunbowale, Elaheh Hadadianpour, Olamide Ishola, Md Majharul Islam, Natalie Ramos, Arvin Saffarian Delkhosh, Elka R. Georgieva

**Author notes:** Equal contribution (Adeyemi Ogunbowale, Elaheh Hadadianpour).

## Abstract

HIV-1 Vpu supports viral adaptation through host-protein interactions. Although mainly membrane-associated, we recently identified a soluble Vpu form that forms a stable complex with Ca²□-bound calmodulin (Ca²□-CaM), potentially influencing Vpu trafficking.

Here, to determine the binding affinity and identify regions of soluble Vpu involved in CaM binding, we used ensemble Förster Resonance Energy Transfer (eFRET). We tested Cy3-labeled full-length (FL) Vpu, a C-terminal fragment (helices 2 and 3), and a Cy3-labeled FL Vpu V22A/W23Y/I33N mutant with substitutions of key residues in Vpu’s helix 1 and helices 1-to-2 loop having a role in the interaction with Ca^2+^-CaM. All Vpu variants were labeled at residue L42C. Ca²□-CaM was tagged with Cy5 at residue S39C. eFRET analysis of 100 nM Cy3-Vpu variants mixed with Cy5-Ca^2+^-CaM (in the range 100 nM–2.5 µM) revealed heterocomplexes formation with dissociation constants (*K_d_*) and binding free energies (Δ*G*). FL Vpu-Ca^2+^-CaM showed highest stability (*K_d_* ∼74 nM, Δ*G* ∼-9.7 kcal/mol), while the truncated C-terminal region and V22A/W23Y/I33N mutant formed weaker complexes with Ca^2+^-CaM (*K_d_* ∼182 nM and 800 nM, Δ*G* ∼-9.2 kcal/mol and ∼-8.3 kcal/mol). The reduced Vpu–Ca²□–CaM stability after disruption of binding sites in and near Vpu helix 1 may explain how this interaction is regulated, including through lipid competition that promotes Vpu membrane insertion. We propose that, at the membrane, hydrophobic helix 1 dissociates from Ca²□–CaM and inserts into the lipid bilayer, weakening the complex and releasing CaM. These findings clarify HIV-1 Vpu interactions with cellular components and may inform antiviral development.

## INTRODUCTION

The human immunodeficiency virus 1 (HIV-1) is the causative agent of HIV/AIDS^1^. HIV-1 employs advanced strategies to enter cells and regulate their functions^2–5^. Understanding HIV-1 pathogenesis requires insight into the virus’s molecular mechanisms within infected cells.

Our focus is on the HIV-1 encoded viral protein U (Vpu), which plays critical role in the virus lifecycle through interacting with cellular membranes and proteins to control their function^6–8^. The protein is expressed in the infected cells and resides and functions in the membranes of trans-Golgi network, ER and plasma membrane^9, 10^. It is one of the smallest HIV-1 proteins with molecular weight of 9-16 kDa, single-pass α-helical type-I transmembrane protein. The amino acid sequence of Vpu consists of a luminal N-terminal domain (residues 1–3), a highly hydrophobic transmembrane domain (residues 4–27, transmembrane helix 1 [TM helix 1]), and a C-terminal cytoplasmic tail encompassing two α-helices (residues 28–81) that are helix 2 and helix 3 interconnected by a linker region (Figure 1)^11, 12^. Through its activity, Vpu alters the integrity and natural abundance of many host proteins at the plasma membrane^13^; it also antagonizes host proteins to prevent premature cell death^14^ and promote efficient viral particle release^15, 16^. Vpu achieves this by exhibiting substantial conformational flexibility, enabling it to interact with a range of host proteins—including the CD4 receptor, tetherin, MHC-I, and MHC-II. These interactions facilitate the degradation, altered localization, or regulatory modification (either down- or up-regulation) of these proteins^1, 9, 17–19^.

**Figure 1.**
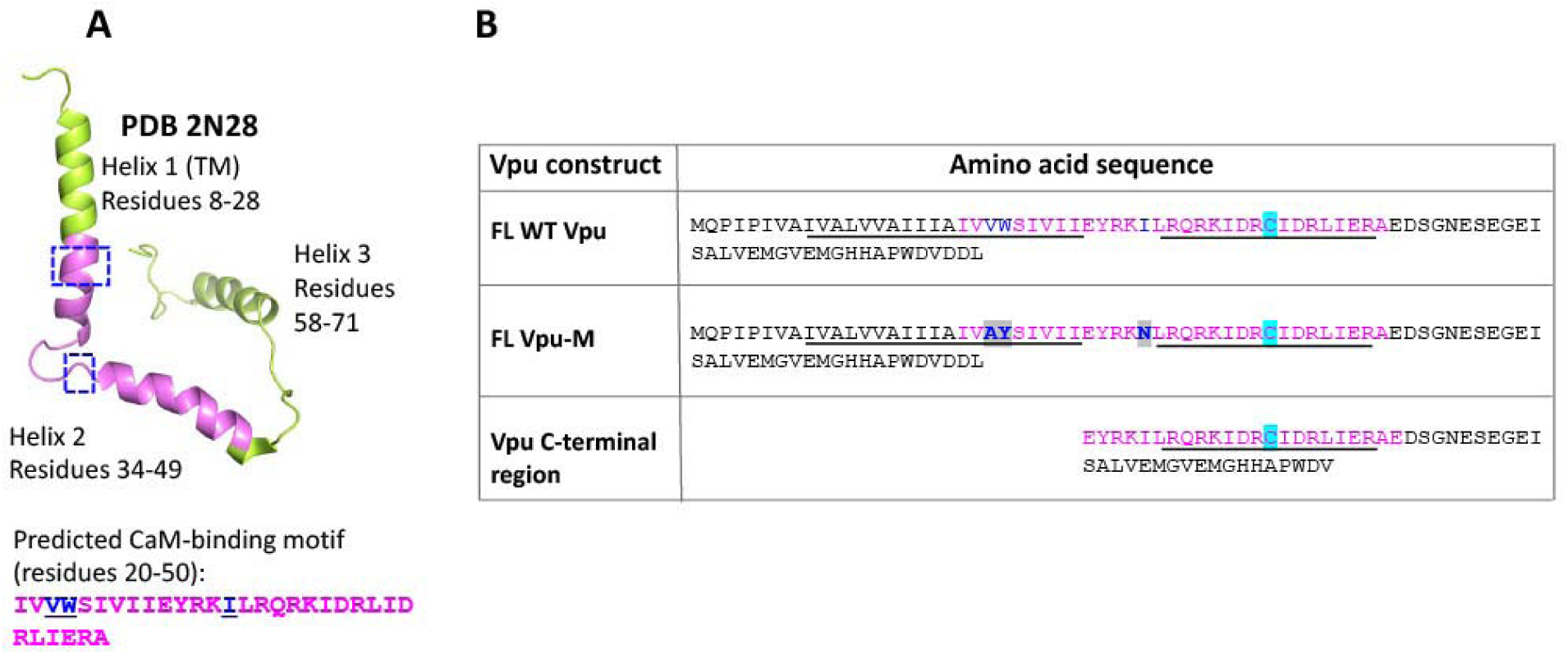
The Vpu protein: (A) The experimentally determined structure of FL Vpu (PDB#2N28) with helices 1, 2, and 3 designated. The earlier predicted CaM-binding motif is in magenta in the structure, and the amino acid sequence of this motif is shown below the structure. The regions of the V22A/W23Y (helix 1) and I33N (helix 1-to-helix 2 loop) mutations are shown in blue boxes; the residues V22, W23, and I33 in the CaM-binding sequence are underlined and in blue. In the shown structure (PDB#2N28 Vpu’s N-terminal region is shorter than the construct we used by 1 amino acid, which reflects the residues’ numbering. (B) The three main Vpu constructs used to test their binding to Ca^2+^-CaM—FL WT Vpu, FL Vpu-M, and truncated Vpu C-terminal region are shown. The helices 1 and 2 are underlined in black. The CaM-binding motif is in magenta. The cysteine residue (L42C mutation) used for labeling with Cy3 donor dye is highlighted in cyan in each construct. The original residues which were substituted for either alanine (A), tyrosine (Y), or asparagine (N) in the V22A/W23Y/I33N mutant are in bold blue and highlighted in gray.

Our lab aims to understand how Vpu interacts with host calmodulin (CaM), which likely influences HIV-1 physiology in infected cells. Elevated CaM levels in HIV-1 infected cells suggest CaM’s key role in the viral adaptation and pathogenesis^20^. Nearly all HIV-1 proteins have predicted non-canonical CaM-binding motifs, and direct interactions of Gag, Nef, Tat, and others with CaM have been reported^21–25^. These associations may facilitate HIV-1 protein trafficking, membrane insertion, regulation of cellular apoptosis, and contribute to CD4+ lymphocyte deterioration^26–32^.

Previously, Vpu was thought to be solely a membrane protein^1, 8, 33, 34^. Recent findings from our lab showed Vpu can also exists in soluble form, forming concentration-dependent homooligomers ^35, 36^. Thus, Vpu resembles other known human and HIV-1 proteins, which undergo physiological transition from soluble to membrane-bound state^37, 38^. We next provided the first experimental evidence that the soluble Vpu interacts with Ca^2+^-bound calmodulin (Ca^2+^-CaM) in equimolar stoichiometry^39^, aligning with earlier CaM-binding sequence predictions in Vpu^25^. The putative CaM-binding motif was believed to span TM helix 1, the loop between helices 1 and 2, and part of helix 2^25^. However, our further amino acid sequence analysis suggested that helix 2 plays a significant role because it contains extensive imperfect stretches of residues found in other proteins having canonical CaM-binding motifs^39–42^. We further found that the C-terminal domain of Vpu (residues 28–82), which includes helices 2 and 3, interacts with Ca^2+^-CaM similarly to full-length Vpu; therefore helix 2 contributes substantially to the interaction with Ca^2+^-CaM^39^. The exact role of soluble Vpu and its interaction with CaM are yet to be characterized in detail. Our current hypothesis is that Vpu utilizes Ca^2+^-CaM for trafficking to the membrane^39^. Furthermore, the Vpu homooligomers could also be viewed as biomacromolecular condensates^43^ having storage or another currently unidentified function, which could be investigated in the future.

Here, we report our results from the quantitative analysis of the association of soluble Vpu with Ca^2+^-CaM. We also examined how different regions in Vpu’s CaM-binding motif contribute to this association. To do so, we used three main constructs of Vpu, which are (i) FL wild-type (WT) Vpu, (ii) FL Vpu with amino acid substitutions V22A, W23Y and I33N in the CaM-binding motif in helix 1 (V22A/W23Y) and the link between helices 1 and 2 (I33N), which we refer to as FL Vpu-M, and (iii) Vpu fragments containing the C-terminal region of WT Vpu (residues 28-82 or 28-78). The FL WT Vpu contained the complete CaM-binding motif (residues 20-50), whereas the second and third Vpu constructs had this motif abrogated either through mutations or truncation (Figure 1). The V22A/W23Y/I33N mutations were carefully selected, so they did not change the hydrophobicity of Vpu helix 1 and charge of the helix 1-helix 2 loop; still, they were sufficiently unlike the native V22/W23/I33 amino acids found in CaM-binding motifs, particularly in the 1-8-14 motif sequences FILVW and ILV ^42^. We utilized ensemble F rster resonance energy transfer (eFRET) to monitor the complex formation between labeled with cyanine 3 (Cy3) Vpu variants and labeled with cyanine 5 (Cy5) FL CaM in the presence of 1 mM CaCl_2_. We found that Vpu interacts reversibly with Ca^2+^-CaM, forming a specific and dynamic complex.

Interestingly, the strength of FL WT Vpu association with Ca^2+^-CaM was higher (*K_d_* ∼ 74 nM) than those for the truncated C-terminal of the WT Vpu (*K_d_* ∼ 182 nM) and FL Vpu-M (*K_d_*∼800 nM), which corresponded to binding free energies (Δ*G*) of ∼ −9.7 kcal/mol, ∼ −9.2 kcal/mol and ∼ −8.3 kcal/mol, respectively. Based on these results, we concluded that both the residues in Vpu’s helix 1 and helix 2 within the CaM-binding motif contribute to the Vpu-Ca^2+^-CaM complex formation. The reduction of ∼0.5-1.4 kcal/mol for Vpu lacking helix 1 or with substitutions in helix 1 suggests this segment of Vpu participated in the heterocomplex stabilization. Although, substantial Vpu-CaM contacts likely occur within Vpu’s helix 2 and the loop between helices 1 and 2, as both Vpu-M and the Vpu C-terminal fragment retained significant binding to Ca^2+^-CaM. These findings may relate to how soluble Vpu uses CaM for membrane trafficking, thus shedding light into an important HIV-1 mechanism. Furthermore, increasing evidence suggests that CaM interactions with proteins from diverse viruses can enhance viral infectivity and spread^44–46^, and our results add to this growing body of work.

## RESULTS

### 1. The Vpu and CaM variants developed and utilized in this study

In this study, we used three main constructs of Vpu (Figure 1B): (i) FL WT Vpu, (ii) Vpu C-terminal region (residues 28-78), and (iii) FL Vpu-M containing the mutations V22A/W23Y/I33N. The FL Vpu and FL Vpu-M were expressed in *E. coli* and purified as SUMO-tagged at their N-terminus proteins (Figure S1). The tag-free FL Vpu and FL Vpu-M were obtained after the SUMO tag removal. To assess the effect of the fusion tag on Vpu-Ca^2+^-CaM binding, we also studied Vpu constructs, which carried the SUMO tag linked to the N-terminus of the Vpu variants, that are SUMO-FL WT Vpu, and SUMO-Vpu C-terminal region (residues 28-82) (Figure S1). These last two constructs are the same as those SUMO-tagged Vpu variant, which we used previously to assess the Vpu-Ca^2+^-CaM complex formation using DEER and fluorescence spectroscopy^39^. All Vpu variants contained a single cysteine mutation, L42C (numbering in FL Vpu), used for fluorescent labeling.

To evaluate the effects of the V22A/W23Y/I33N mutations in FL Vpu, we predicted the structure of this Vpu variant using AlphaFold^47^. No alterations were predicted in the helical region surrounding these amino acids (Figure S2), suggesting that these mutations are likely to have only a minimal, if any, effect on Vpu structure. This outcome is expected, as V22 and W23 were deliberately substituted with A and Y, respectively, which possess properties like those of the original residues at these positions; the I33N mutation was selected to maintain the properties of the helix 1-to-helix 2 loop. Nevertheless, the combination of V22A, W23Y, and I33N substitutions, which are located within the CaM-binding motif in Vpu, weakened Vpu-Ca^2+^-CaM association.

The CaM protein was the same as the one we used earlier with a single cysteine residue at position S39C^39^, which was used for labeling with Cy5 acceptor dye.

### 2. Vpu self-associates above concentrations of 100 nM affecting binding to Ca^2+^-CaM

As we observed earlier, the truncated C-terminal fragment of tag-free Vpu forms homo-oligomers at concentrations as low as 500 nM, and FL Vpu also self-associates into soluble oligomers^35, 36, 39^. To assess the extent to which Vpu homooligomerization influences the analysis of Vpu-Ca^2+^-CaM complex formation, we examined the binding of FL Vpu and the Vpu C-terminal fragment to Ca^2+^-CaM. In series of experiments, the concentrations of the Cy3-labeled Vpu constructs (Cy3-Vpu) varied from 100 nM to 850 nM, while the Cy5-labeled Ca^2+^-CaM (Cy5-Ca^2+^-CaM) concentration was maintained constant at 500 nM. Fluorescence spectra for all combinations of Cy3-Vpu and Cy5-Ca^2+^-CaM were collected over the 555–800 nm range (Figure 3, left panels, Figures S3 and S4). Characteristic Cy3 emission peaks were observed at approximately 570 nm and 610 nm, accompanied by a Cy5 acceptor emission peak near 675 nm upon formation of the Cy3-Vpu variant–Cy5-Ca^2+^-CaM complex. The FRET efficiency (*E_FRET_*) for each sample was calculated using the equation^48^:

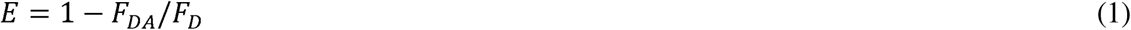

Where *E* is the FRET efficiency (*E_FRET_*), *F_D_* is the fluorescence intensity of the Cy3 donor at the maximum at about 610 nm without Cy5-Ca^2+^-CaM in the sample; and *F_DA_* is the fluorescence intensity at the same peak maximum for each sample in the presence of Cy5-Ca^2+^-CaM.

This method for *E_FRET_*determination is based on the donor quenching approach.

Maximum *E_FRET_* was observed at 100 nM for the FL WT and truncated C-terminal region of Vpu. At higher concentrations of Vpu variants, *E_FRET_* decreased, particularly for the truncated Vpu C-terminal fragment (Figure 2, right panels). These results indicate that, above 100 nM, Vpu homooligomerization reduces the pool of monomers available for Ca^2+^-CaM binding, thereby weakening the interaction. This finding is consistent with our previous DEER spectroscopy results obtained at micromolar protein concentrations, where pronounced self-association of Vpu variants likely limited Ca^2+^-CaM binding to approximately 50%, despite the presence of excess CaM^39^. Moreover, these results strongly support our hypothesis that Vpu variants form an equimolar complex with Ca^2+^-CaM, which is in agreement with the data showing that the amplitude of the DEER signal (modulation depth) for doubly spin-labeled Ca^2+^-CaM does not increase upon binding to Vpu (i.e., no higher order oligomers were formed) while the Vpu homooligomers disassemble upon addition of Ca^2+^-CaM^39^.

**Figure 2.**
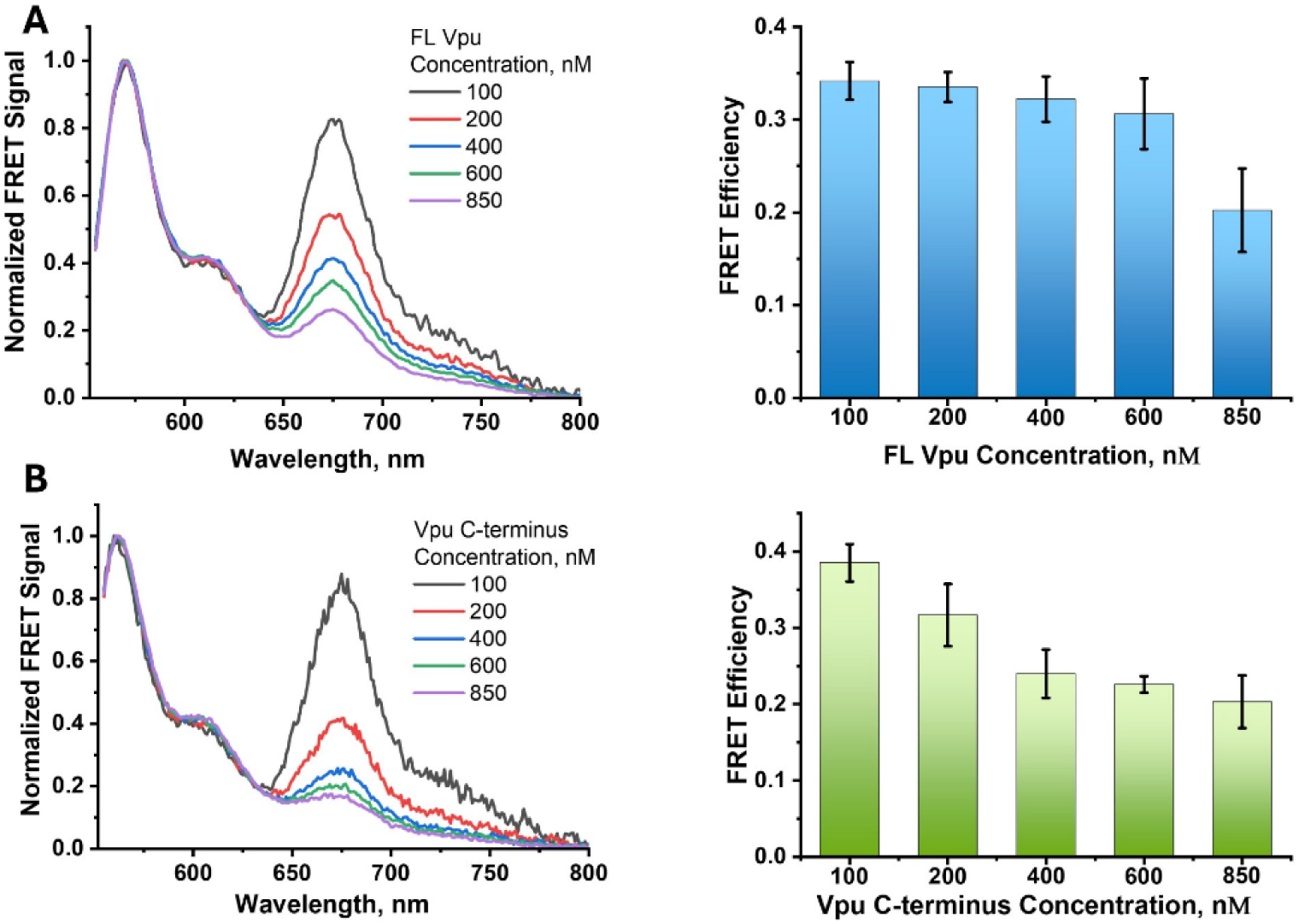
Normalized eFRET spectra and FRET efficiencies of the interaction between Vpu variants and Ca^2+^-CaM. eFRET spectra (left) were normalized to unity at the maximum Cy3 emission to facilitate comparison of the relative Cy5 acceptor emission across samples. Cy3-labeled Vpu variants were measured at increasing concentrations (100 nM – 850 nM) in the presence of a constant concentration of 500 nM Cy5-Ca^2+^-CaM. Full-length (FL) Vpu data are shown in the upper panels (A), and data for the Vpu C-terminal region are shown in the lower panels (B). Normalization highlights the progressive decrease in Cy5 emission relative to Cy3 emission as the concentration of Cy3-Vpu increases. FRET efficiencies, *E_FRET_* (right), were calculated from the corresponding raw, non-normalized eFRET data and are presented as mean ± SD. The decrease in *E_FRET_* with increasing Vpu concentration is consistent with reduced Vpu-CaM association at higher Vpu concentrations, likely resulting from increased Vpu homo-oligomerization.

Based on these results, we used 100 nM in the subsequent experiments to determine the dissociation constant *K_d_* and binding free energy Δ*G* of the Vpu-Ca^2+^-CaM heterocomplex, as described below.

### 3. Vpu forms a reversible, dynamic complex with Ca^2+^-CaM

To evaluate the reversibility of complex formation, we performed competition experiments using Cy3-FL WT Vpu and the Cy3-Vpu C-terminal region with either Cy5-Ca²□-CaM alone or mixtures containing Cy5-Ca²□-CaM and unlabeled Ca²□-CaM. Cy3-labeled Vpu variants were maintained at 100 nM, whereas Cy5-Ca²□-CaM was used at 200 nM or 450 nM. After formation of the Cy3-Vpu–Cy5-Ca²□-CaM complex, unlabeled Ca²□-CaM was added at final concentrations of 0.7 µM or 5 µM. In all cases, the addition of unlabeled Ca²□-CaM reduced FRET efficiency and Cy5-Ca^2+^-CaM emission intensity (Figures 3 and S5). This effect was detectable at 0.7 µM, where *E_FRET_*decreased by approximately 10–25%, and was more pronounced at 5 µM, where *E_FRET_*reductions of 65–75% were observed. These results indicate that unlabeled Ca²□-CaM competitively displaced Cy5-Ca²□-CaM from their complexes with Vpu variants. Thus, after the addition of non-labeled Ca^2+^-CaM, a new steady state containing protein monomers and complexes of Cy3-Vpu variants with either Cy5-Ca^2+^-CaM or non-labeled Ca^2+^-CaM was established.

**Figure 3.**
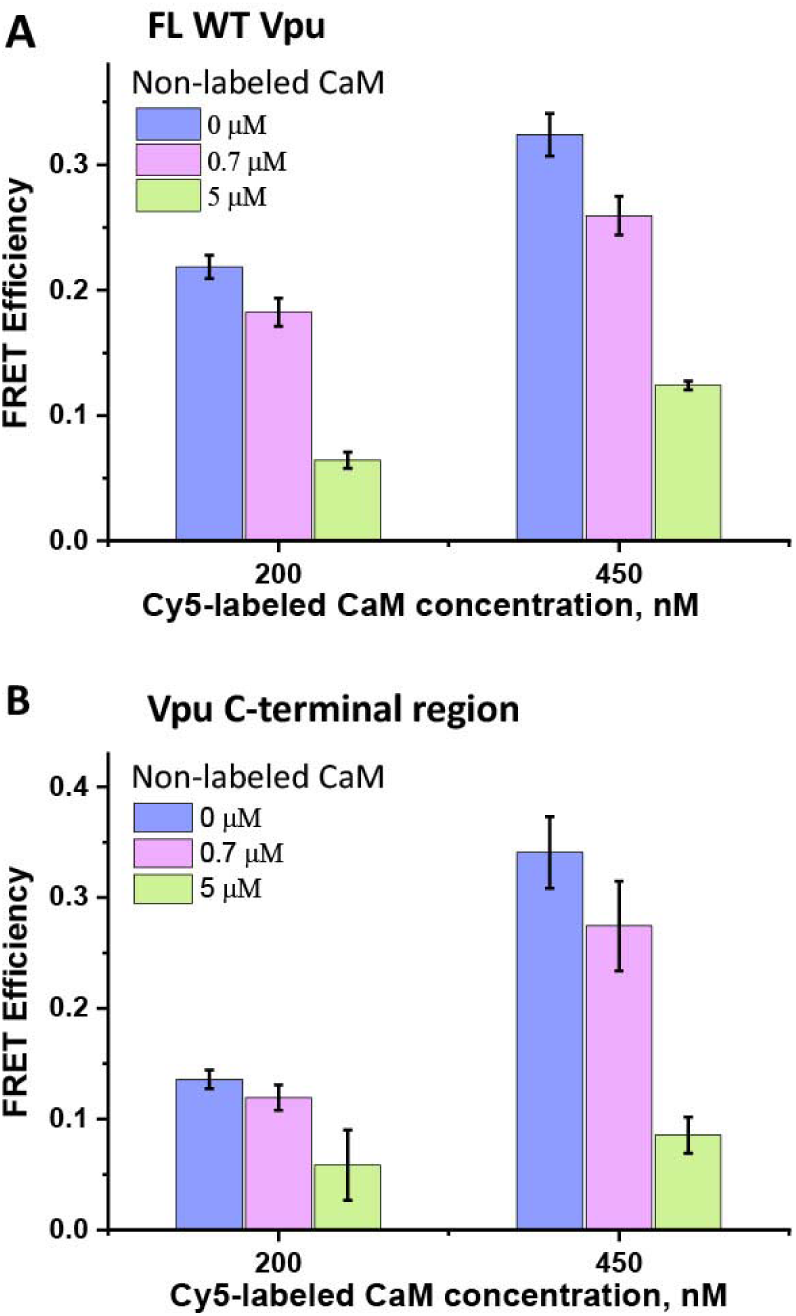
FRET efficiencies for FL WT Vpu-Ca^2+^-CaM (A) and Vpu C-terminal fragment-Ca^2+^-CaM (B) complex formation. In all cases, constant concentration of Cy3-Vpu variants at 100 nM was used. Two concentrations of Cy5-Ca^2+^-CaM that are 200 nM and 450 nM were used. Additionally, to some of the samples, 0.7 µM and 5 µM of non-labeled Ca^2+^-CaM were added. The results for different sample conditions are color coded as indicated in the figure. A clear decrease in *E_FRET_*was observed after the addition of non-labeled Ca^2+^-CaM. The mean data and standard deviations of three experiments at each condition are shown.

These competition experiments demonstrate that Vpu-Ca²□-CaM complex formation is reversible and governed by dynamic equilibrium rather than nonspecific aggregation. The displacement of Cy5-Ca²□-CaM by excess unlabeled Ca²□-CaM validates the use of equilibrium-based binding analysis and supports the biological plausibility of CaM acting as a temporary binding partner for soluble Vpu.

### 4. All studied Vpu constructs bind Ca^2+^-CaM with nanomolar-range dissociation constants and free energies of ∼ – 9.7 kcal/mol to – 8.3 kcal/mol

Here, we examined how distinct regions of Vpu’s CaM-binding motif contribute to the stability of the Vpu-Ca^2+^-CaM complex.

We used eFRET to monitor binding between Cy3-labeled Vpu variants and Cy5-labeled Ca^2+^-CaM. The analysis included FL WT Vpu, FL Vpu-M, and the Vpu C-terminal region. We also compared SUMO-tagged Vpu variants with their corresponding tag-free forms to assess whether the N-terminal SUMO tag affected Ca^2+^-CaM binding. All Vpu variants were at constant concentration of 100 nM, and series of samples with increasing Ca^2+^-CaM concentrations in the range from 100 nM to 2.5 µM were analyzed. The fluorescence spectra of Cy3-Vpu variants at 100 nM alone were also recorded as the donor-only controls for comparison with the spectra obtained in the presence of increasing concentrations of Cy5-Ca^2+^-CaM. As expected, as the Cy5-Ca^2+^-CaM concentration increased, the intensity of the Cy3-Vpu fluorescence peaks with maxima at ∼570 nm and 610 nm decreased proportionally to the Vpu variant-Ca^2+^-CaM complex formation due to Cy3-to-Cy5 energy transfer. Simultaneously, the intensity of Cy5-Ca^2+^-CaM emission with a maximum at ∼675 nm also increased, reflecting the heterocomplex formation (Figures 4A, 5A, 6A, and Figures S6 and S7).

**Figure 4.**
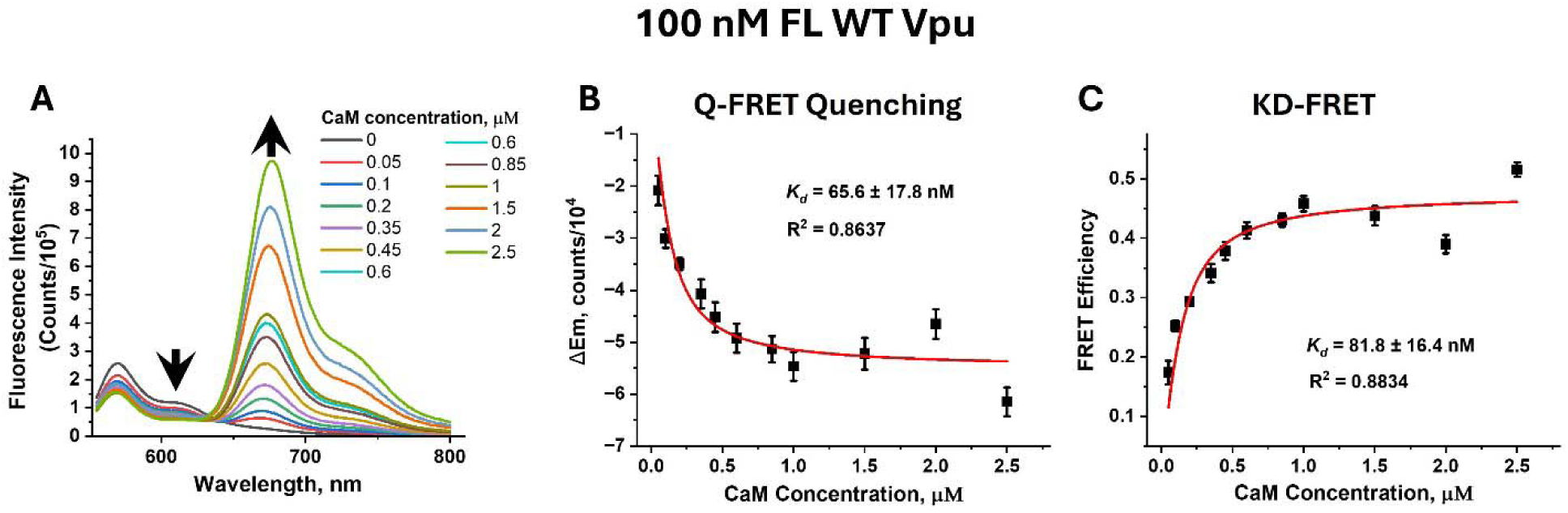
eFRET data and analyses for the interaction of 100 nM Cy3-FLWT Vpu with Cy5-Ca^2+^-CaM at increasing concentration in the range from 0 to 2.5 µM. The original eFRET data are shown in (A)—The Cy3 emission decreases while Cy5 emission increases upon Cy3-Vpu-Ca2+-CaM complex formation. The background-corrected eFRET spectra (in A) were used to estimate *K_d_*. The results from fitting the Cy3 donor fluorescence decay at ∼ 609 nm, shown with the arrow, (method of Quantitative FRET Quenching), and fitting the FRET efficiencies (*E_FRET_*) (KD-FRET) to obtain *K_d_*-s are shown in (B) and (C), respectively. The estimated *K_d_*-s and quality of data fits are listed in (B) and (C). There is a very good agreement between the *K_d_* values derived using the two methods. All data are averages of triplicate samples. In (B) and (C), the mean values and standard deviations of three experiments at each condition are shown.

**Figure 5.**
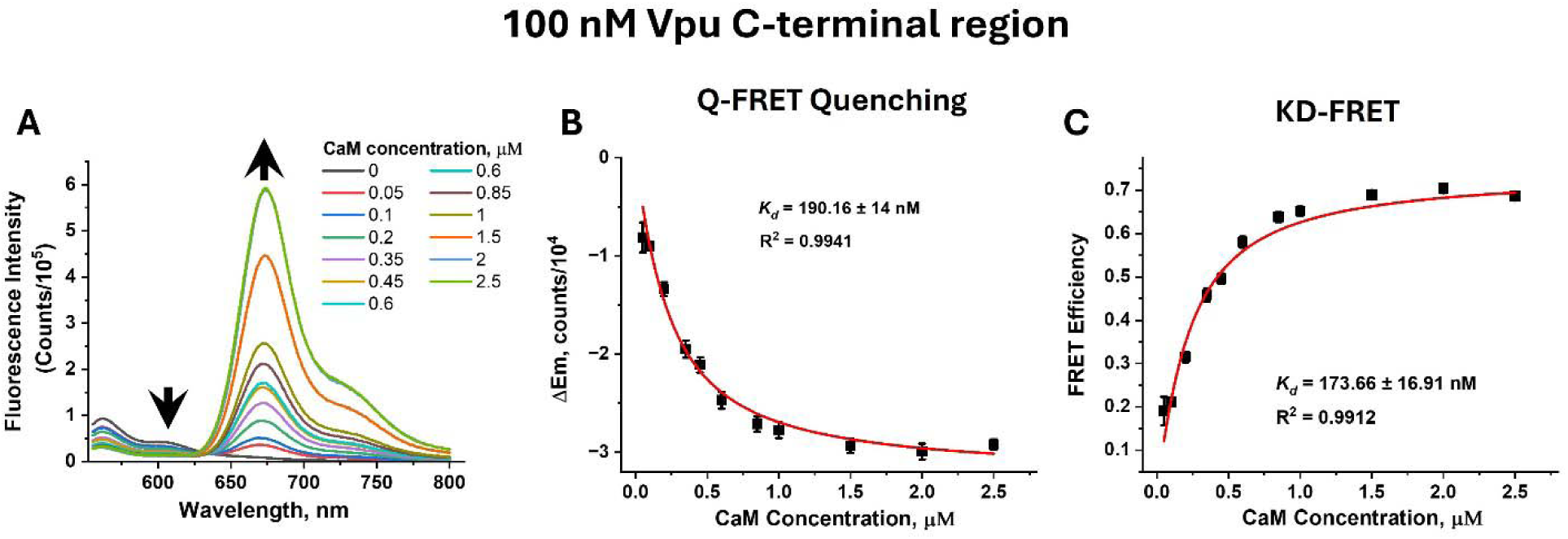
eFRET data and analyses for the interaction of 100 nM Cy3-Vpu C-terminal region with Cy5-Ca2+-CaM at increasing concentrations in the range from 0 to 2.5 µM. The original eFRET data are shown in (A)—The Cy3 emission decreases while Cy5 emission increases upon Cy3-Vpu-Ca2+-CaM complex formation. The background-corrected eFRET spectra (in A) were used to estimate Kd. The results from fitting the Cy3 donor fluorescence decay at ∼ 602 nm, shown with the arrow (method of Quantitative FRET Quenching), and fitting the FRET efficiencies (EFRET) (KD-FRET) to obtain *K_d_*-s are shown in (B) and (C), respectively. The estimated *K_d_*-s and quality of data fits are shown in (B) and (C) inset. There is a very good agreement between the *K_d_*values derived using the two methods. All data are averages of triplicate samples. In (B) and (C), the mean values and standard deviations of three experiments at each condition are shown.

**Figure 6.**
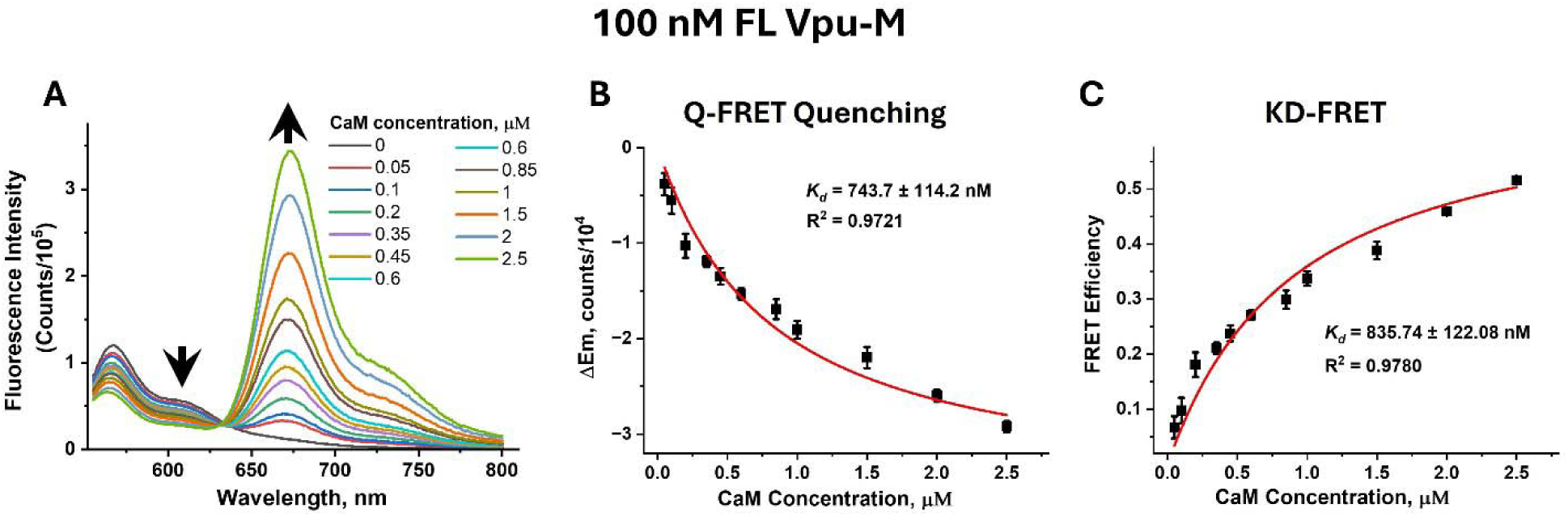
eFRET data and analyses for the interaction of 100 nM Cy3-FLVpu-M with Cy5-Ca^2+^-CaM at increasing concentrations in the range from 0 to 2.5 µM. The original eFRET data are shown in (A)—The Cy3 emission decreases while Cy5 emission increases upon Cy3-Vpu-Ca2+-CaM complex formation. The background-corrected eFRET data (in A) were used to estimate *K_d_*. The results from fitting the Cy3 donor fluorescence decay at ∼ 606 nm, shown with the arrow (method of Quantitative FRET Quenching), and fitting the FRET efficiencies (*E_FRET_*) (KD-FRET) to obtain *K_d_*-s are shown in (B) and (C), respectively. The estimated *K_d_*-s and quality of data fits are shown in (B) and (C) inset. There is a very good agreement between the *K_d_* values derived using the two methods. All data are averages of triplicate samples. In (B) and (C), the mean values and standard deviations of three experiments at each condition are shown.

We then used these eFRET data sets to estimate the dissociation constants (*K_d_*-s) and binding free energies (Δ*G*-s) of the WT Vpu variants with Ca^2+^-CaM. To do so, we plotted the change in the Cy3-Vpu variant intensity of the peak at ∼603-609 nm (coinciding with Cy5 absorption) vs. Cy5-Ca^2+^-CaM concentration (Figures 4B, 5B, and 6B). Thereafter, adopting the method of quantitative FRET quenching^49^ we estimated the *K_d_*-s for the complexes of Vpu variants with Ca^2+^-CaM by fitting the data to the equation:

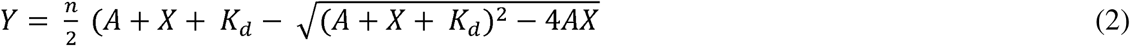

Where, n is a constant related to FRET efficiency between donor (Vpu construct) and acceptor (CaM) – obtained from fittings, A is a constant concentration of donor (Cy3-Vpu), and X is the varying concentration of acceptor (Cy5-CaM).

Further, we employed the FRET-based method (KD-FRET), which was used to quantify protein– protein interactions in bacterial cells, relying on the *E*_FRET_ ^50^. This method is also based on donor quenching approach. For each series of samples containing 100 nM Vpu variant and Cy5-Ca^2+^-CaM concentrations in the range of 100 nM to 2.5 µM, the values of *E_FRET_*were calculated using Eq. (1) and plotted against the concentrations of Cy5-Ca^2+^-CaM (Figures 4C, 5C, and 6C). The *K_d_*-s were obtained by fitting these data to the equation:

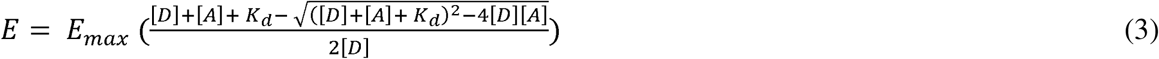

Where *E_max_*is the maximum FRET efficiency (obtained from fittings), [D] is the concentration of donor labeled protein (Vpu), and [A] is the varying concentration of acceptor labeled protein (CaM) We used the obtained *K_d_-*s to calculate the Δ*G*-s of each of the Vpu-Ca^2+^-CaM complexes. Both quantitative FRET quenching and KD-FRET analyses of the same eFRET data sets yielded very close *K_d_* and Δ*G* values (Figures 4, 5, and 6).

#### 4.1. FL WT Vpu binds Ca^2+^-CaM with relatively high affinity

The conducted binding analyses of Cy3-FL WT Vpu with Cy5-Ca2+-CaM yielded *K_d_* values of 65.6 ± 17.8 nM by quantitative FRET quenching and 81.8 ± 16.4 nM by KD-FRET (Figure 4), indicating that FL Vpu binds Ca^2+^-CaM with relatively high affinity. The calculated Δ*G* was approximately −9.7 kcal/mol, consistent with formation of a stable protein-protein complex^51, 52^. Further, these binding parameters suggest that the Vpu-Ca^2+^-CaM complex lies near the lower limit of stability relative to CaM complexes with host proteins, which typically exhibit Δ*G* values ranging from −10 to −15 kcal/mol^53^. However, the results for Vpu agree with prior findings about HIV-1 MA protein-Ca^2+^-CaM complexes, which reported a *K_d_* of about 37 nm for FL CaM-binding motif in this protein ^54^. Together, these findings suggest that the association of HIV-1 proteins with Ca^2+^-CaM may support more dynamic cycles of complex formation and dissociation.

#### 4.2. The truncated C-terminal region of Vpu has reduced affinity to Ca^2+^-CaM

The calculated *K_d_* for the Vpu C-terminal region-Ca^2+^-CaM complex increased to 190.16 ± 14 nM (quantitative FRET quenching) and 173.7 ± 16.9 nM (KD-FRET) (Figure 5), which is about 2.5 times higher than the value for the FL Vpu. The Δ*G* was to ∼ −9.2 kcal/mol, a reduction by less than 1 kcal/mol compared to FL Vpu. The slight increase in *K_d_* and reduction in Δ*G* suggest small but measurable decrease in the binding affinity of the Vpu C-terminal region to Ca^2+^-CaM compared to FL Vpu. Nevertheless, the observed binding was still substantial. It is worth noting that the Vpu C-terminal fragment (Figure 1B) contained the residues forming the helix-1-to-helix 2 loop and the complete soluble C-terminal region of Vpu, including helix 2, thus comprising both positively charged and hydrophobic residues found to play roles in the binding of host proteins to Ca^2+^-CaM^25^. Therefore, apparently, this Vpu region has a key role in the association with Ca^2+^-CaM.

We further noted that the *E_FRET_* for the Cy3-FL Vpu was close to 2 times lower compared to that of Cy3-Vpu C-terminal fragment under the same concentrations of Cy5-Ca^2+^-CaM (Figures 4C and 5C). This was a persistent result despite the lower labeling efficiency of the Vpu C-terminal region compared to FL Vpu (Materials and Methods). This could be due to steric effects imposed by the FL Vpu or altered conformations of the proteins in the FL-Vpu- and Vpu C-terminus-Ca^2+^-CaM complexes, which results in an increased distance between the Cy3 donor and Cy5 acceptor in the FL Vpu-Ca^2+^-CaM complex^55^.

#### 4.3. FL Vpu carrying the V22A/W23Y/I33N mutations has reduced affinity to Ca^2+^-CaM

It was predicted that both hydrophobic (mostly located in helix 1) and positively charged residues (in the loop between helices 1 and 2 and in helix 2) in the Vpu polypeptide contribute to the interaction with CaM^25, 39^. In addition, in the current work, we found that the truncated C-terminal region of Vpu has slightly reduced binding affinity to Ca^2+^-CaM compared to FL WT Vpu. This prompted us to study the contribution of the hydrophobic residues in Vpu’s helix 1 and helix 1-to-helix 2 loop to the Vpu-Ca^2+^-CaM complex stabilization. To test this, we engineered a triple mutant (FL Vpu-M) with substitutions V22A/W23Y of residues in the imperfect 1-8-14 CaM-binding motif encompassing the residues IVVWS in Vpu’s helix 1, and an additional mutation I33N toward helix 2 but still in the proximity of helix 1 (Figure 1B). We conducted the same eFRET-based assay and data analysis, as for the other two Vpu variants (Figure 6). Strikingly, for this Vpu-M, we observed further decrease in its affinity to Ca^2+^-CaM, as the *K_d_* increased to ∼800 nM and Δ*G* decreased to ∼ −8.3 kcal/mol, yielding a difference of 1.4 kcal/mol compared to FL WT Vpu. Thus, these results suggest that the residues V22/W23/I33 indeed contribute measurably to the Vpu-Ca^2+^-CaM interaction. The observed shift in *K_d_* and Δ*G* is in the direction of those caused by the truncation of Vpu’s helix 1, therefore supporting the involvement of Vpu’s IVVWS motif and hydrophobic contacts in the Vpu-Ca^2+^-CaM complex.

#### 4.4. Soluble FL WT Vpu forms a specific and stable complex with Ca^2+^-CaM, while mutations in the Vpu CaM-binding motif in the region of helix 1 and the helix 1-helix 2 loop as well as the complete truncation of helix 1 destabilize this complex

The *K_d_* and Δ*G* values for the interaction of all soluble Vpu constructs used in this study with Ca^2+^-CaM are summarized in Tables 1 and 2 and Figure 7. As mentioned above, the Δ*G* values for all complexes suggest relatively stable specific complexes, as they fell within the range of −7 kcal/mol to −20 kcal/mol reported for other protein-protein complexes^52^. However, the Vpu-Ca^2+^-CaM affinity and complex stability decreases upon the substitution of key hydrophobic amino acids V22A/W23Y/I33N in Vpu’s helix 1 and the following loop, as well as complete removal of Vpu’s helix 1. Both modifications are in the highly hydrophobic Vpu’s helix1 and its vicinity, which is known to associate with membrane adopting transmembrane location^18, 33, 56^. Thus, the observed results may be linked to Vpu’s transition from soluble to membrane-bound state and release of CaM, but future studies will be needed to test this hypothesis.

**Figure 7.**
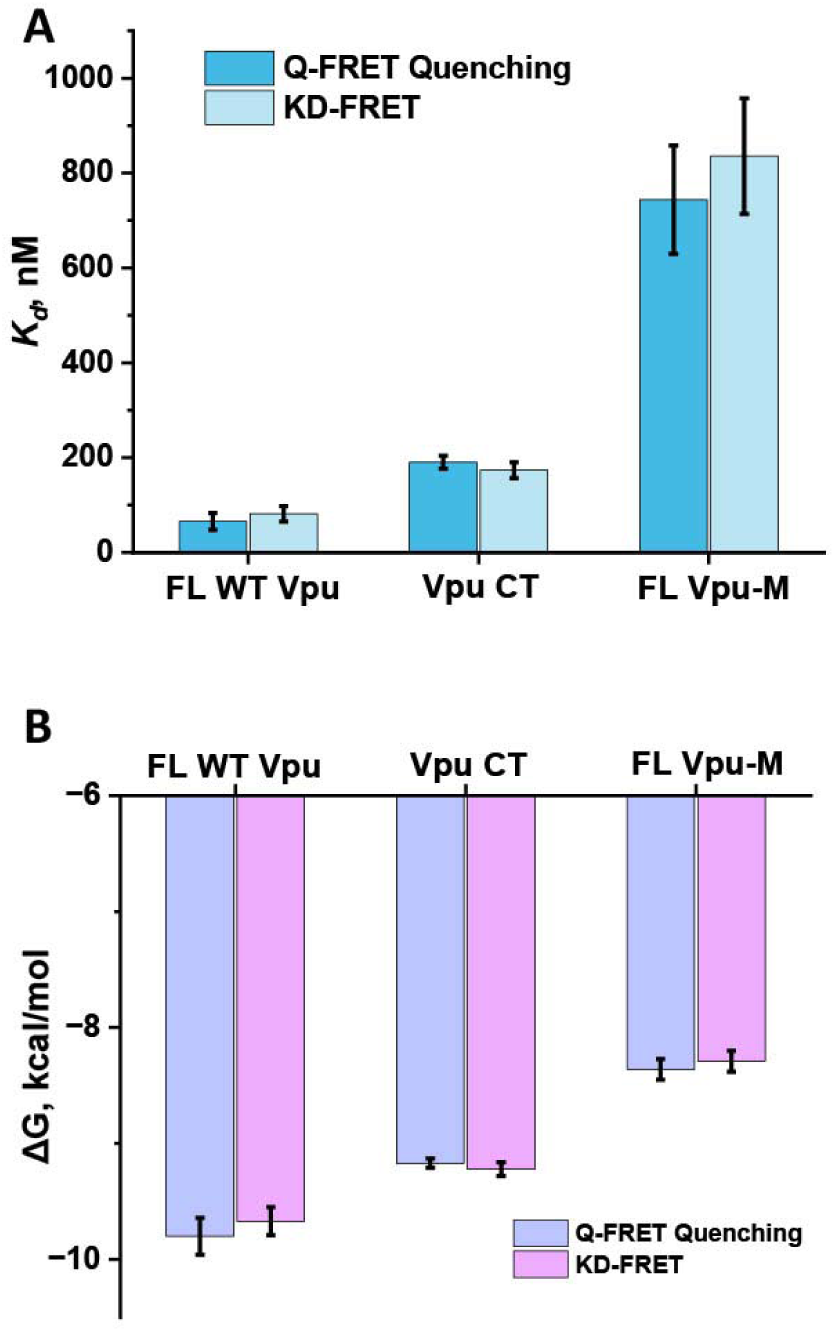
Bar-diagrams plots of the dissociation constants (*K_d_*) (A) and binding free energies (Δ*G*) (B) values for the complexes of Vpu constructs to Ca^2+^-CaM. The data obtained using both the Quantitative FRET Quenching (Q-FRET Quenching) and KD-FRET methods are shown. The mean values and standard deviations of three experiments at each condition are shown.

**Table 1.** The estimated *K_d_*values for the complexes of all Vpu constructs used in this study with Ca^2+^-CaM. The mean values ±standard deviation (SD) are shown. The data obtained using Q-FRET Quenching and KD-FRET are shown.

| Vpu construct | $K_d \pm SD$, nM<br>Q-FRET<br>Quenching | $K_d \pm SD$, nM<br>KD-FRET |
| --- | --- | --- |
| FL WT Vpu | $65.6 \pm 17.8$ | $81.8 \pm 16.4$ |
| Vpu C-terminal region | $190.16 \pm 14$ | $173.66 \pm 16.91$ |
| FL Vpu-M | $743.7 \pm 114.2$ | $835.74 \pm 122.1$ |

**Table 2.** The calculated binding energies ΔG-s for the complexes of all Vpu constructs used in this study with Ca^2+^-CaM. The mean values ±standard deviation (SD) are shown. The data obtained using Q-FRET Quenching and KD-FRET are shown.

| Vpu construct | $\Delta G \pm SD$, kcal/mol<br>Q-FRET<br>Quenching | $\Delta G \pm SD$, kcal/mol<br>KD-FRET |
| --- | --- | --- |
| FL WT Vpu | $-9.80 \pm 0.16$ | $-9.67 \pm 0.12$ |
| Vpu C-terminal region | $-9.17 \pm 0.04$ | $-9.22 \pm 0.06$ |
| FL Vpu-M | $-8.36 \pm 0.09$ | $-8.29 \pm 0.09$ |

#### 4.5. The SUMO tag in the SUMO-FL WT Vpu and SUMO-Vpu C-terminal region has very small effect of the binding with Ca^2+^-CaM

Next, we aimed to determine the effect of SUMO tag on the Vpu-Ca^2+^-CaM binding affinities. This is an important question to address, as we used SUMO-tagged Vpu variants in our preceding study^39^ and others used SUMO-tagged proteins prior to us^57^. Therefore, we determined the *K_d_* and Δ*G* for the association of SUMO-FL WT Vpu and SUMO-Vpu C-terminal with Ca^2+^-CaM. We analyzed the eFRET data for these complexes (Figures S6 and S7) in the same way we did for SUMO tag-free Vpu variants, yielding *K_d_*-s of ∼140 nM and ∼561 nM and Δ*G*-s of ∼ −9.3 kcal/mol and ∼ −8.5 kcal/mol, respectively. Thus, the presence of SUMO tag indeed reduced the binding but by only about 0.67 kcal/mol or less. Interestingly, the effect of SUMO tag on binding was more pronounced for the truncated fragment of Vpu vs. FL Vpu. The reason likely is that in the FL Vpu the SUMO tag is fused to the N-terminal and therefore, it is further away from the CaM-binding motif in Vpu, compared to Vpu C-terminal where SUMO is near the binding site.

## DISCUSSION

In our previous studies, we uncovered that the assumed exclusively transmembrane HIV-1 Vpu protein can exist in a soluble form^35, 36^ and later we revealed that the soluble Vpu forms an equimolar complex with Ca^2+^-CaM^39^ which we believe is linked to Vpu’s trafficking to the membrane site. Indeed, the interactions of HIV-1 proteins seem to be critical for HIV-1 physiology in the infected cells. Besides Vpu, other HIV-1 proteins, e.g., the MA domain of Gag, Nef, Tat, etc., associate with CaM, and it is thought these interactions aid HIV-1 proteins’ trafficking, insertion into the membrane, regulation of cellular apoptosis, and contribute to the CD4+-lymphocytes deterioration^26–32^. Notably, it was found that, compared to non-infected cells, the HIV-1 infected cells have higher CaM levels, particularly in the membrane-bound organelle subcellular fraction^20^, suggesting that CaM may be used to traffic the synthesized in a soluble form HIV-1 proteins to cellular locations, including membranes, where they are active. These findings support the possible significant role of CaM in the HIV-1 life cycle. Additionally, CaM was linked to the replication and infectivity of other viruses—e.g., it interacts with Ebola matrix protein VP40^58^, highlighting the broader role of viral protein-CaM interactions.

Here, we aimed to gain deeper insight into how the HIV-1 Vpu protein binds with Ca^2+^-CaM. Our previous work showed that Vpu variants form homooligomers^35, 36, 39^, so we performed experiments to investigate the dissociation of these complexes. We found that at a concentration of 100 nM, all Vpu variants were mainly present in their monomeric form (Figure 2). This led us to use this concentration in binding assays between Vpu variants and Ca^2+^-CaM, under the assumption that only monomeric forms contribute to heterocomplex kinetics. This assumption was supported by the partial Vpu-Ca^2+^-CaM complexes formation at micromolar concentration observed by DEER^39^—At levels above 100 nM, fewer Vpu monomers are available as more join homooligomers. Thus, our findings further support that Vpu variants interact with Ca^2+^-CaM as monomers in an equimolar ratio, in agreement with our prior DEER data^39^.

Because of the observed homooligomerization of soluble Vpu in our prior^35, 36, 39^ and current studies, it is worth discussing what could be the origin of these oligomers. Besides, homooligomerization of the soluble form of other HIV-1 proteins that are directed to and function in the cellular membranes has also been observed. For example, it was found that the HIV-1 Nef protein forms homodimers and homotrimers in solution^3^. Another study found that the HIV-1 MA alone forms trimer or higher order oligomer in solution^59^. However, despite it having been found that Nef and MA interact directly with Ca^2+^-CaM ^22, 28^, no indication that CaM-binds to either Nef or MA oligomer was reported. On the contrary, results from protein sedimentation and size exclusion chromatography (SEC) suggested that MA and CaM form a 1:1 complex^29^. Therefore, the physiological significance of these HIV-1 proteins, i.e., Vpu, Nef and MA, homooligomers is not well understood. One possibility is that they are easily formed when in an isolated state *in vitro*; however, in the presence of a binding partner, e.g., Ca^2+^-CaM, their homooligomerization may be irrelevant or a transient state. One possibility could be that the soluble oligomers of HIV-1 proteins are biomacromolecular condensates formed to separate from the aqueous environment; they serve as a storage of these proteins when they do not engage in protein-protein or protein lipid interactions or could represent an uncharacterized functional state. Indeed, membraneless biomolecular condensates of HIV-1 proteins have been studied and discussed in the literature^60^. The behavior of HIV-1 proteins thus adds to the much broader idea about protein clustering and liquid-liquid phase separations, which has been a topic of extensive research in the recent years, as it is linked to both physiological and disease processes^61, 62^.

The competitive eFRET binding assays conducted using Cy3-Vpu, Cy5-Ca^2+^-CaM, and excess unlabeled Ca^2+^-CaM confirmed that the Vpu-Ca^2+^-CaM interaction is reversible and dynamic, supporting its physiological relevance.

Using eFRET measurements across a Ca^2+^-CaM concentration range of 100 nM to 2.5 µM, we determined the *K_d_*-s for Vpu variants containing the full-length CaM-binding motif or modified versions with amino acid substitutions or N-terminal truncation.

We found that FL WT Vpu forms a relatively stable complex with *K_d_* of ∼74 nM and Δ*G* of ∼-9.7 kcal/mol, while its truncated C-terminal region (*K_d_* ∼182 nM, Δ*G* ∼-9.2 kcal/mol) and FL Vpu-M with V22A/W23Y/I33N mutations (*Kd* ∼800 nM, Δ*G* ∼-8.3 kcal/mol) bind less tightly. These results indicate that the hydrophobic contacts in and proximal to Vpu’s helix 1 contribute to the stability of Vpu-Ca^2+^-CaM association. This may relate to Vpu’s transition from soluble to membrane-bound states, requiring helix 1 to unbind Ca^2+^-CaM following insertion in the membrane. Thereafter, CaM is released as the complex destabilizes. This is plausible, as it was found earlier that the HIV-1 MA protein as well has two modes of interaction with Ca^2+^-CaM that include the FL CaM-binding amino acid sequence (residues 8–43), but shorter regions of residues 11-28 and 31-46 also bind with reduced affinity^54^, which might be a mechanism used by HIV-1 proteins.

We further observed that the binding free energies of the Vpu-Ca^2+^-CaM complexes, although consistent with specific Ca^2+^-CaM–protein interactions, fall within the strong-to-moderate range reported for CaM complexes with host proteins^51–53^. This finding may reflect an evolutionarily refined mechanism by which HIV-1 engages host proteins, enabling the resulting protein-protein complexes to remain sufficiently dynamic to support efficient complex formation and dissociation.

Our findings contribute to the understanding of how HIV-1 proteins interact with host CaM. The quantitative data which we provide will be useful in directing drug design to regulate the Vpu-host protein (CaM) interactions. The *K_d_* and Δ*G* values for the Vpu-Ca^2+^-CaM complex provide a foundation for identifying drugs with stronger binding, as previously demonstrated ^63, 64^.

## MATERIALS AND METHODS

### 1. Protein designs, cloning, mutagenesis, expression, and purification

The DNA-s encoding the SUMO-FL Vpu and SUMO-C-terminal region of Vpu with histidine tags were commercially synthesized and cloned in pET15b vector (GenScript, Inc.), as described^39^. They contained the L42C mutation for labeling with a Cy3 donor. The peptide encompassing the Vpu residues 28-78 with a cysteine residue at position L42C (numbering in FL Vpu) was commercially synthesized (RS Synthesis). The CaM mutant S39C was produced as previously described^39^. The SUMO-FL Vpu-M containing the V22A/W23Y/I33N mutations were generated using site-directed mutagenesis (GenScript, Inc.). The expression and purification procedure are fully described in Ishola et al. 2026^39^.

### 2. Removing the SUMO tag

The SUMO tag was removed through thrombin digestion: The Nickel affinity–purified Vpu variants in a buffer of 20 mM Tris pH 8.0, 100 mM NaCl, 5% (w/v) glycerol, 1 mM DDM, and 100 μM TCEP and 10 mM CaCl was mixed with human thrombin (Millipore Sigma) at a ratio of 20 U/mg thrombin/1 mg Vpu protein. The reaction was allowed to proceed overnight at 22°C with gentle rotation. On the next day, the reaction mixture was incubated with Ni²-NTA resin pre-equilibrated in binding buffer containing 20 mM Tris pH 7.4, 150 mM NaCl, 1 mM CaCl, 5% (w/v) glycerol, 1 mM β-DDM, and 100 μM TCEP. Binding was performed for 1 hour at 4°C with gentle mixing. Thereafter, the flow-through containing the SUMO tag and thrombin was discarded; the resin with bound Vpu protein was washed with 5 resin volumes of the same buffer. Next, the Vpu variant was eluted using 320 mM Imidazole. Afterwards, the Imidazole was removed from the Vpu proteins, and they were concentrated in centrifuge concentrators with 3 kDa MWCO at 4 °C. The high degree of thrombin removal was confirmed using SDS-PAGE and Western Blotting (WB).

### 3. Labeling of the proteins with Cy3 donor and Cy5 acceptor

Initially, the β-DDM was removed from the FL WT Vpu and FL Vpu-M (with and without SUMO tag) by washing the proteins with a buffer containing 20 mM Tris pH 7.4, 150 mM NaCl, 1 mM CaCl_2_, 50 µM TCEP, and 5% glycerol. The Vpu C-terminal region (with and without SUMO tag) was also in this buffer. Buffer exchange and β-DDM removal was performed using several dilution and concentration steps at 4 °C to ensure complete replacement of the initial buffers. The final concentrations of all protein constructs were determined using a NanoDrop^TM^ One spectrophotometer.

Thereafter, all these buffer-exchanged proteins were labeled with the cysteine-specific donor cyanine3-maleimide (Cy3) at a 1:5 protein-to-dye molar ratio and incubated for 3.5 h at 22 °C under constant agitation. Thereafter, the samples were placed at 4 °C and then incubated overnight under constant agitation. During the incubation, the samples were wrapped in aluminum foil to protect the fluorophores from light exposure, ensuring labeling occurred in a dark environment. On the next day, the unreacted Cy3 was removed by passing the protein/dyes mixtures 2 times through the NAP 5 column (Cytiva). The Cy3-labeled Vpu variants were in a final buffer of 20 mM Tris pH 7.4, 150 mM NaCl, 1 mM CaCl_2_, 50 µM TCEP, and 5% glycerol, which was used in all FRET experiments.

Ca^2+^-CaM was labeled using the same protocol but with the cysteine-specific cyanine5-maleimide (Cy5) acceptor instead of Cy3 donor, using the previously described protocol^39^.

The details of the procedure for handling the Vpu C-terminal peptide were described in detail in Ishola et al. 2026 ^39^.

### 4. Quantification of Cy3/Cy5 labeling efficiency

The degree of labeling (labeling efficiency) is the molar ratio of bound dye concentration to protein concentration after removal of unreacted dye ^65^. The dye and protein concentrations were quantified using the protein and labels function of the NanoDrop^TM^ One spectrophotometer. The parameters for Cy3-maleimide and Cy5-maleimide were entered into the system, and the calculations were performed as described by Chedda et al. (2023)^65^. For Cy3-maleimide, dye concentration was determined from A555 using an extinction coefficient of 150,000 M ¹ cm ¹. For Cy5-maleimide, dye concentration was determined from A646 using an extinction coefficient of 250,000 M ¹ cm ¹. Protein concentration was calculated from the absorbance at 280 nm after correcting for dye absorbance at 280 nm using the dye-specific correction factors (CF280 = 0.09 for Cy3 and 0.04 for Cy5). Thus, protein and dye concentrations were calculated according to:

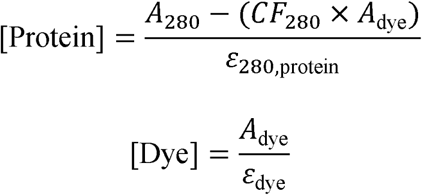

Labeling efficiency was then expressed as the molar dye-to-protein ratio:

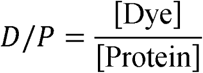

For the Cy3-labeled Vpu C-terminal peptide, the protein extinction coefficient at 280 nm was ε_280,protein_ 8604M□¹ cm□¹; therefore, the working equations were:

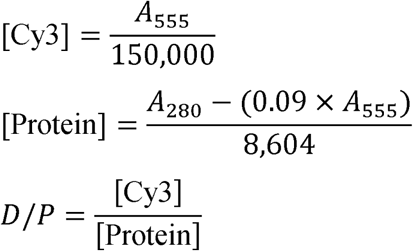

Accordingly, the labeling efficiencies reported in Table 1 represent mol dye per mol protein rather than an arbitrary instrument-generated ratio.

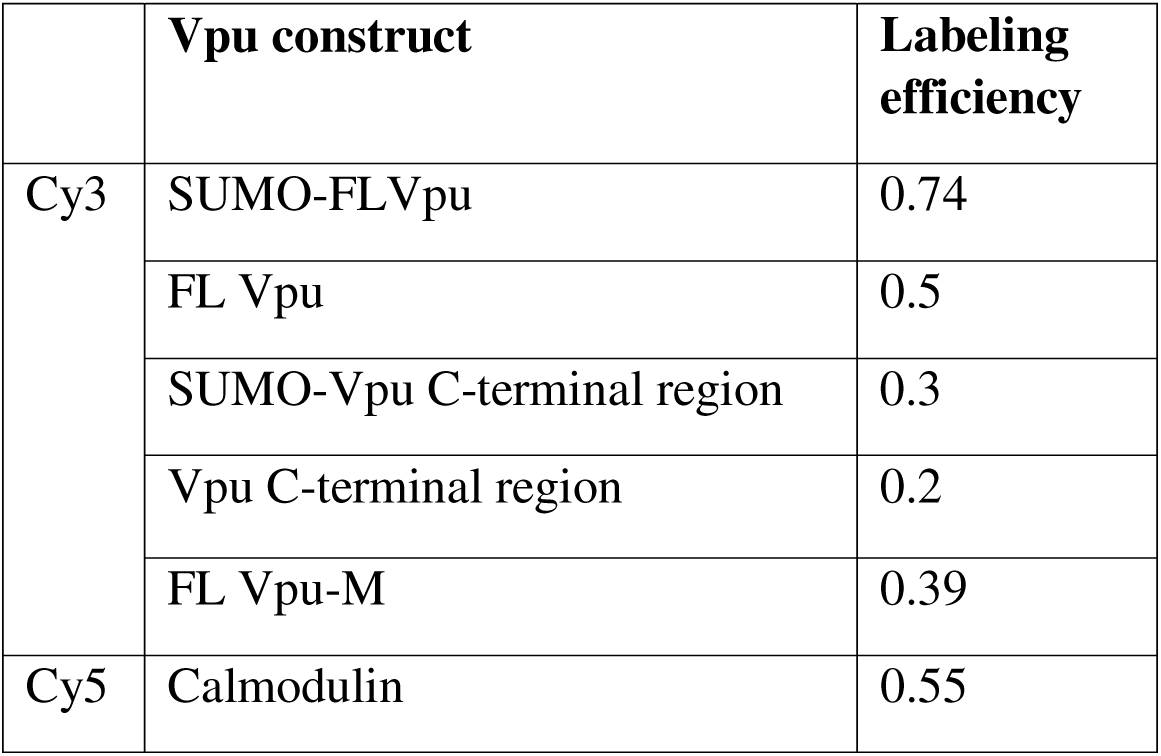

|  | **Vpu construct** | **Labeling efficiency** |
| --- | --- | --- |
| Cy3 | SUMO-FLVpu | 0.74 |
|  | FL Vpu | 0.5 |
|  | SUMO-Vpu C-terminal region | 0.3 |
|  | Vpu C-terminal region | 0.2 |
|  | FL Vpu-M | 0.39 |
| Cy5 | Calmodulin | 0.55 |

### 5. Ensemble FRET experiments and data analysis

For these experiments, triplicate samples (three independently prepared and measured replicates) were used to evaluate the reproducibility of the measurements. The data points shown in Figures 2–5 represent the mean values and standard deviations derived from these triplicate measurements. The mixing time between labeled Vpu and labeled Ca² –CaM was tightly controlled, and all samples within each series were measured under identical conditions. This ensured that potential artifacts such as donor emission bleaching or dilution effects were minimized, preventing false eFRET signals.

All eFRET data were collected at 25 °C using a temperature-controlled FS5 spectrofluorometer (Edinburgh Instruments).

The following samples were prepared and eFRET experiments conducted:

a. To determine the optimal range of Cy5-Vpu variant concentration for quantitative analysis, samples containing increasing concentration (100, 200, 400, 600, and 850 nM) of tag-free Cy3-labeled FL Vpu and Vpu C-terminal region mixed with constant 500 nM Cy5-labeled CaM. The FRET data were collected using an FS5 spectrofluorometer (Edinburgh Instrument) with the donor excitation at 550 nm (1 nm excitation and 2 nm emission bandwidth), and emission recorded from 555 to 800 nm.
b. To evaluate the dynamic nature of the Vpu-Ca²-CaM complexes, Cy3-labeled Vpu constructs, including FL WT Vpu and the Vpu C-terminal region, were maintained at a constant concentration of 100 nM, whereas Cy5-labeled Ca²-CaM was used at concentrations of 200 nM or 450 nM. In selected samples, unlabeled Ca²-CaM was subsequently added at final concentrations of either 0.7 µM or 5 µM. All samples were adjusted to a final volume of 1000 µL to maintain constant concentrations of the labeled proteins, with only the concentration of unlabeled Ca²-CaM varied. Thus, the samples were prepared by initially combining 800 µL of FRET buffer (20 mM Tris pH 7.4, 150 mM NaCl, 1 mM CaCl_2_, 50 µM TCEP) with the appropriate volumes of Cy3-labeled Vpu constructs and Cy5-labeled Ca²-CaM, followed by a 10 min incubation in a dark room at room temperature (RT) to allow complex formation. Unlabeled Ca²-CaM was then added for each concentration condition, and the buffer (20 mM Tris pH 7.4, 150 mM NaCl, 1 mM CaCl_2_, 50 µM TCEP) was used to bring the final volume to 1000 µL. The mixtures were incubated for an additional 5 min in dark at RT before FRET measurements were acquired. Fluorescence spectra were collected using Cy3 donor excitation at 550 nm, with 2 nm excitation and emission bandwidths, and emission was recorded from 555 to 800 nm.
c. To conduct quantitative analysis of Vpu variant-Ca^2+^-CaM binding, samples containing a mixture of 100 nM SUMO-FL Vpu, FL Vpu, SUMO-Vpu C-terminal fragment, Vpu C-terminal fragment, FL Vpu-M were mixed with increasing concentration of Ca^2+^-CaM at 0.05 µM, 0.1 µM, 0.2 µM, 0.35 µM, 0.45 µM, 0.6 µM, 0.85 µM, 1 µM, 1.5 µM, 2 µM, and 2.5 µM. The FRET data were collected using an FS5 spectrofluorometer (Edinburgh Instrument) with the donor excitation at 550 nm (3 nm excitation and emission bandwidth), and emission recorded from 555 to 800 nm. Each sample was prepared by mixing Cy3-Vpu variant with Cy5-Ca^2+^-CaM in the FRET buffer containing 1 mM CaCl_2_ in a cuvette. The protein mixtures were incubated at RT in a dark room for 10 minutes to allow protein interaction.

The dissociation constant (*K_d_*) for the Vpu-variant–Ca² –CaM complexes were determined using two complementary FRET-based approaches. In the first method, Quantitative FRET Quenching, the donor fluorescence was monitored as the concentration of the acceptor was increased during the titration, and the extent of donor quenching (ΔEm) was calculated relative to the donor-only control to quantify the fraction of bound complex. ΔEm at the donor maximum wavelength was plotted against the total CaM concentration and fitted using the non-linear regression model described by Jiang et al. ^66^. In this model (equation 2), *n* is a fitting parameter related to the FRET efficiency between the Cy3-labeled Vpu donor and Cy5-labeled CaM acceptor; *A* is the fixed donor concentration; *Y* represents the difference in donor emission intensity in the presence and absence of acceptor; and *X* is the varying acceptor concentration. The second method, KD-FRET, involved calculating FRET efficiency using the donor-quenching equation from Shresta et al.^67^ and subsequently fitting the efficiency values to a quadratic binding model (equation 1) as described by Yi et al., 2025 ^68^. In this model, Emax denotes the maximum FRET efficiency obtained from curve fitting, while [D] and [A] correspond to the concentrations of donor-labeled Vpu variant and acceptor-labeled CaM, respectively. Together, these models provide complementary ways of extracting dissociation constants from donor quenching behavior, with all *K_d_*values calculated from triplicate measurements and reported as mean ± standard deviation. The R^2^ fitting goodness factor for all cases was equal or greater than 0.86. All the graph plotting and fitting to determine the *K_d_* values were performed using Origin Pro 2023 (Origin Lab, Northampton, MA).

## AUTHOR CONTRIBUTION

AO—experimental design, experiment, data analysis and interpretation, figures, writing the experimental procedures; EH— experimental design, experiment, data analysis and interpretation, figures, writing the experimental procedures; MMI—experiment; NR— experiment; ASD—experiment, data analysis and interpretation; ERG—conception, experimental design, data analysis and interpretation, figures, writing the manuscript, supervision, finds acquisition.

## Supporting information

Manuscript

## ACKNOWLEDGEMENTS

This work was supported by the Gilead Research Scholars in HIV Award to ERG. We thank Juan Camilo Rueda Amador for helping with initial experiments. ERG thanks Dr. Peter Borbat for fruitful discussions that led to the conception of this work. We thank the Reviewer for the valuable suggestions and comments that helped us to improve the quality of this work.

## CONFLICT OF INTERESTS

The authors declare no conflict of interest.

## DATA AVAILABILITY

Nearly all experimental data produced through this study are shown in the manuscript. The data can be further obtained by contacting the corresponding author with a reasonable request.

## Notes

### Competing Interest Statement

The authors have declared no competing interest.

### Summary of Updates

In the revised manuscript we: 1. Modified the title to better reflect the work. 2. Included new data and conclusions. 3. Significantly restructured and rewrote the manuscript to include the new data and conclusions. 4. Most of the figures were revised. 5. Supplementary file was expanded substantially to include most of the raw data.

## References

(1) Khan, N.; Geiger, J. D. Role of Viral Protein U (Vpu) in HIV-1 Infection and Pathogenesis. Viruses 2021, 13 (8). DOI: 10.3390/v13081466 From NLM Medline.

(2) Altfeld, M.; Gale, M., Jr. Innate immunity against HIV-1 infection. Nat Immunol 2015, 16 (6), 554–562. DOI: 10.1038/ni.3157 From NLM Medline.

(3) Arold, S.; Hoh, F.; Domergue, S.; Birck, C.; Delsuc, M. A.; Jullien, M.; Dumas, C. Characterization and molecular basis of the oligomeric structure of HIV-1 nef protein. Protein Sci 2000, 9 (6), 1137–1148. DOI: 10.1110/ps.9.6.1137 From NLM Medline.

(4) Hladik, F.; McElrath, M. J. Setting the stage: host invasion by HIV. Nat Rev Immunol 2008, 8(6), 447–457. DOI: 10.1038/nri2302 From NLM Medline.

(5) Ramdas, P.; Sahu, A. K.; Mishra, T.; Bhardwaj, V.; Chande, A. From Entry to Egress: Strategic Exploitation of the Cellular Processes by HIV-1. Front Microbiol 2020, 11, 559792. DOI: 10.3389/fmicb.2020.559792 From NLM PubMed-not-MEDLINE.

(6) Dube, M.; Bego, M. G.; Paquay, C.; Cohen, E. A. Modulation of HIV-1-host interaction: role of the Vpu accessory protein. Retrovirology 2010, 7, 114. DOI: 10.1186/1742-4690-7-114 From NLM Medline.

(7) Langer, S.; Hammer, C.; Hopfensperger, K.; Klein, L.; Hotter, D.; De Jesus, P. D.; Herbert, K. M.; Pache, L.; Smith, N.; van der Merwe, J. A.;, et al. HIV-1 Vpu is a potent transcriptional suppressor of NF-kappaB-elicited antiviral immune responses. Elife 2019, 8. DOI: 10.7554/eLife.41930.

(8) Strebel, K.; Klimkait, T.; Martin, M. A. A novel gene of HIV-1, vpu, and its 16-kilodalton product. Science 1988, 241 (4870), 1221–1223. DOI: 10.1126/science.3261888.

(9) Gonzalez, M. E. Vpu Protein: The Viroporin Encoded by HIV-1. Viruses 2015, 7 (8), 4352– 4368. DOI: 10.3390/v7082824 From NLM Medline.

(10) Hussain, A.; Das, S. R.; Tanwar, C.; Jameel, S. Oligomerization of the human immunodeficiency virus type 1 (HIV-1) Vpu protein--a genetic, biochemical and biophysical analysis. Virol J 2007, 4, 81. DOI: 10.1186/1743-422X-4-81.

(11) Cohen, E. A.; Terwilliger, E. F.; Sodroski, J. G.; Haseltine, W. A. Identification of a protein encoded by the vpu gene of HIV-1. Nature 1988, 334 (6182), 532–534.

(12) Strebel, K.; Klimkait, T.; Martin, M. A. A novel gene of HIV-1, vpu, and its 16-kilodalton product. Science 1988, 241 (4870), 1221–1223.

(13) Cong, L.; Sugden, S. M.; Leclair, P.; Lim, C. J.; Pham, T. N.; Cohen, É. A. HIV-1 Vpu promotes phagocytosis of infected CD4+ T cells by macrophages through downregulation of CD47. Mbio 2021, 12 (4), e01920–01921.

(14) Wildum, S.; Schindler, M.; Mu nch, J.; Kirchhoff, F. Contribution of Vpu, Env, and Nef to CD4 down-modulation and resistance of human immunodeficiency virus type 1-infected T cells to superinfection. Journal of virology 2006, 80 (16), 8047–8059.

(15) Neil, S. J.; Zang, T.; Bieniasz, P. D. Tetherin inhibits retrovirus release and is antagonized by HIV-1 Vpu. Nature 2008, 451 (7177), 425–430.

(16) Van Damme, N.; Goff, D.; Katsura, C.; Jorgenson, R. L.; Mitchell, R.; Johnson, M. C.; Stephens, E. B.; Guatelli, J. The interferon-induced protein BST-2 restricts HIV-1 release and is downregulated from the cell surface by the viral Vpu protein. Cell host & microbe 2008, 3 (4), 245–252.

(17) Willey, R.; Maldarelli, F.; Martin, M. A.; Strebel, K. Human immunodeficiency virus type 1 Vpu protein induces rapid degradation of CD4. Journal of virology 1992, 66 (12), 7193–7200.

(18) González, M. E. Vpu protein: The viroporin encoded by HIV-1. Viruses 2015, 7 (8), 4352–4368.

(19) McNatt, M. W.; Zang, T.; Bieniasz, P. D. Vpu binds directly to tetherin and displaces it from nascent virions. PLoS Pathog 2013, 9 (4), e1003299. DOI: 10.1371/journal.ppat.1003299 From NLM Medline.

(20) Radding, W.; Pan, Z. Q.; Hunter, E.; Johnston, P.; Williams, J. P.; McDonald, J. M. Expression of HIV-1 envelope glycoprotein alters cellular calmodulin. Biochem Biophys Res Commun 1996, 218 (1), 192–197. DOI: 10.1006/bbrc.1996.0034 From NLM Medline.

(21) Alaimo, A.; Alberdi, A.; Gomis-Perez, C.; Fernandez-Orth, J.; Gomez-Posada, J. C.; Areso, P.; Villarroel, A. Cooperativity between calmodulin-binding sites in Kv7.2 channels. J Cell Sci 2013, 126 (Pt 1), 244–253. DOI: 10.1242/jcs.114082 From NLM Medline.

(22) Hayashi, N.; Matsubara, M.; Jinbo, Y.; Titani, K.; Izumi, Y.; Matsushima, N. Nef of HIV-1 interacts directly with calcium-bound calmodulin. Protein Sci 2002, 11 (3), 529–537. DOI: 10.1110/ps.23702 From NLM Medline.

(23) Kurokawa, H.; Osawa, M.; Kurihara, H.; Katayama, N.; Tokumitsu, H.; Swindells, M. B.; Kainosho, M.; Ikura, M. Target-induced conformational adaptation of calmodulin revealed by the crystal structure of a complex with nematode Ca(2+)/calmodulin-dependent kinase kinase peptide. J Mol Biol 2001, 312 (1), 59–68. DOI: 10.1006/jmbi.2001.4822 From NLM Medline.

(24) O’Neil, K. T.; DeGrado, W. F. How calmodulin binds its targets: sequence independent recognition of amphiphilic alpha-helices. Trends Biochem Sci 1990, 15 (2), 59–64. DOI: 10.1016/0968-0004(90)90177-d From NLM Medline.

(25) McQueen, P.; Donald, L. J.; Vo, T. N.; Nguyen, D. H.; Griffiths, H.; Shojania, S.; Standing, K. G.; O’Neil, J. D. Tat peptide-calmodulin binding studies and bioinformatics of HIV-1 protein-calmodulin interactions. Proteins 2011, 79 (7), 2233–2246. DOI: 10.1002/prot.23048 From NLM Medline.

(26) Miller, M. A.; Mietzner, T. A.; Cloyd, M. W.; Robey, W. G.; Montelaro, R. C. Identification of a calmodulin-binding and inhibitory peptide domain in the HIV-1 transmembrane glycoprotein. AIDS Res Hum Retroviruses 1993, 9 (11), 1057–1066. DOI: 10.1089/aid.1993.9.1057 From NLM Medline.

(27) Sham, S. W.; McDonald, J. M.; Micoli, K. J.; Krishna, N. R. Solution structure of a calmodulin-binding domain in the carboxy-terminal region of HIV type 1 gp160. AIDS Res Hum Retroviruses 2008, 24 (4), 607–616. DOI: 10.1089/aid.2007.0202 From NLM Medline.

(28) Radding, W.; Williams, J. P.; McKenna, M. A.; Tummala, R.; Hunter, E.; Tytler, E. M.; McDonald, J. M. Calmodulin and HIV type 1: interactions with Gag and Gag products. AIDS Res Hum Retroviruses 2000, 16 (15), 1519–1525. DOI: 10.1089/088922200750006047 From NLM Medline.

(29) Ghanam, R. H.; Fernandez, T. F.; Fledderman, E. L.; Saad, J. S. Binding of calmodulin to the HIV-1 matrix protein triggers myristate exposure. J Biol Chem 2010, 285 (53), 41911– 41920. DOI: 10.1074/jbc.M110.179093 From NLM Medline.

(30) Vlach, J.; Samal, A. B.; Saad, J. S. Solution structure of calmodulin bound to the binding domain of the HIV-1 matrix protein. J Biol Chem 2014, 289 (12), 8697–8705. DOI: 10.1074/jbc.M113.543694 From NLM Medline.

(31) Dick, A.; Cocklin, S. Subtype Differences in the Interaction of HIV-1 Matrix with Calmodulin: Implications for Biological Functions. Biomolecules 2021, 11 (9). DOI: 10.3390/biom11091294 From NLM Medline.

(32) Pan, G.; Zhou, T.; Radding, W.; Saag, M. S.; Mountz, J. D.; McDonald, J. M. Calmodulin antagonists inhibit apoptosis of CD4+ T-cells from patients with AIDS. Immunopharmacology 1998, 40 (2), 91–103. DOI: 10.1016/s0162-3109(98)00018-6 From NLM Medline.

(33) Sharpe, S.; Yau, W. M.; Tycko, R. Structure and dynamics of the HIV-1 Vpu transmembrane domain revealed by solid-state NMR with magic-angle spinning. Biochemistry 2006, 45 (3), 918–933. DOI: 10.1021/bi051766k From NLM Medline.

(34) Zhang, H.; Lin, E. C.; Das, B. B.; Tian, Y.; Opella, S. J. Structural determination of virus protein U from HIV-1 by NMR in membrane environments. Biochim Biophys Acta 2015, 1848 (11 Pt A), 3007–3018. DOI: 10.1016/j.bbamem.2015.09.008 From NLM Medline.

(35) Majeed, S.; Adetuyi, O.; Borbat, P. P.; Majharul Islam, M.; Ishola, O.; Zhao, B.; Georgieva, E. R. Insights into the oligomeric structure of the HIV-1 Vpu protein. J Struct Biol 2023, 215 (1), 107943. DOI: 10.1016/j.jsb.2023.107943 From NLM Medline.

(36) Majeed, S.; Dang, L.; Islam, M. M.; Ishola, O.; Borbat, P. P.; Ludtke, S. J.; Georgieva, E. R. HIV-1 Vpu protein forms stable oligomers in aqueous solution via its transmembrane domain self-association. Sci Rep 2023, 13 (1), 14691. DOI: 10.1038/s41598-023-41873-0 From NLM Medline.

(37) Hsu, Y. T.; Wolter, K. G.; Youle, R. J. Cytosol-to-membrane redistribution of Bax and Bcl-X(L) during apoptosis. Proc Natl Acad Sci U S A 1997, 94 (8), 3668–3672. DOI: 10.1073/pnas.94.8.3668 From NLM Medline.

(38) Ciftci, H.; Tateishi, H.; Koiwai, K.; Koga, R.; Anraku, K.; Monde, K.; Dag, C.; Destan, E.; Yuksel, B.; Ayan, E.;, et al. Structural insight into host plasma membrane association and assembly of HIV-1 matrix protein. Sci Rep 2021, 11 (1), 15819. DOI: 10.1038/s41598-021-95236-8 From NLM Medline.

(39) Ishola, O.; Islam, M. M.; Hadadianpour, E.; Borbat, P. P.; Ogunbowale, A.; Amador, J. C. R.; Georgieva, E. R. The soluble state of the HIV-1 Vpu protein forms a complex with Ca2+-calmodulin. Protein Science 2026, 35, e70487. DOI: doi: 10.1002/pro.70487Digital.

(40) O’Day, D. H.; Huber, R. J. Calmodulin binding proteins and neuroinflammation in multiple neurodegenerative diseases. BMC Neurosci 2022, 23 (1), 10. DOI: 10.1186/s12868-022-00695-y From NLM Medline.

(41) Osawa, M.; Tokumitsu, H.; Swindells, M. B.; Kurihara, H.; Orita, M.; Shibanuma, T.; Furuya, T.; Ikura, M. A novel target recognition revealed by calmodulin in complex with Ca2+-calmodulin-dependent kinase kinase. Nat Struct Biol 1999, 6 (9), 819–824. DOI: 10.1038/12271 From NLM Medline.

(42) Rhoads, A. R.; Friedberg, F. Sequence motifs for calmodulin recognition. FASEB J 1997, 11 (5), 331–340. DOI: 10.1096/fasebj.11.5.9141499 From NLM Medline.

(43) Sagan, S. M.; Weber, S. C. Let’s phase it: viruses are master architects of biomolecular condensates. Trends Biochem Sci 2023, 48 (3), 229–243. DOI: 10.1016/j.tibs.2022.09.008 From NLM Medline.

(44) Buresova, K.; Nesporova, T.; Prchal, J.; Banerjee, S.; Castoralova, M.; Hodbodova, L.; Kukacka, Z.; Junkova, P.; Ruml, T. Structural and functional insights into calmodulin-mediated lipid binding and proteolytic cleavage of the M-PMV matrix protein. J Biol Chem 2026, 302 (2), 111102. DOI: 10.1016/j.jbc.2025.111102 From NLM Medline.

(45) Chattopadhyay, S.; Basak, T.; Nayak, M. K.; Bhardwaj, G.; Mukherjee, A.; Bhowmick, R.; Sengupta, S.; Chakrabarti, O.; Chatterjee, N. S.; Chawla-Sarkar, M. Identification of cellular calcium binding protein calmodulin as a regulator of rotavirus A infection during comparative proteomic study. PLoS One 2013, 8 (2), e56655. DOI: 10.1371/journal.pone.0056655 From NLM Medline.

(46) Wang, Y.; Gong, Q.; Wu, Y.; Huang, F.; Ismayil, A.; Zhang, D.; Li, H.; Gu, H.; Ludman, M.; Fatyol, K.;, et al. A calmodulin-binding transcription factor links calcium signaling to antiviral RNAi defense in plants. Cell Host Microbe 2021, 29 (9), 1393–1406 e1397. DOI: 10.1016/j.chom.2021.07.003 From NLM Medline.

(47) Jumper, J.; Evans, R.; Pritzel, A.; Green, T.; Figurnov, M.; Ronneberger, O.; Tunyasuvunakool, K.; Bates, R.; Zidek, A.; Potapenko, A.;, et al. Highly accurate protein structure prediction with AlphaFold. Nature 2021, 596 (7873), 583–589. DOI: 10.1038/s41586-021-03819-2 From NLM Medline.

(48) Shrestha, D.; Jenei, A.; Nagy, P.; Vereb, G.; Szollosi, J. Understanding FRET as a research tool for cellular studies. Int J Mol Sci 2015, 16 (4), 6718–6756. DOI: 10.3390/ijms16046718 From NLM Medline.

(49) Jiang, L.; Xiong, Z.; Song, Y.; Lu, Y.; Chen, Y.; Schultz, J. S.; Li, J.; Liao, J. Protein-Protein Affinity Determination by Quantitative FRET Quenching. Sci Rep 2019, 9 (1), 2050. DOI: 10.1038/s41598-018-35535-9 From NLM Medline.

(50) Yi, S.; Kim, E.; Yang, S.; Kim, G.; Bae, D. W.; Son, S. Y.; Jeong, B. G.; Ji, J. S.; Lee, H. H.; Hahn, J. S.;, et al. Direct Quantification of Protein-Protein Interactions in Living Bacterial Cells. Adv Sci (Weinh) 2025, 12 (19), e2414777. DOI: 10.1002/advs.202414777 From NLM Medline.

(51) Day, E. S.; Cote, S. M.; Whitty, A. Binding efficiency of protein-protein complexes. Biochemistry 2012, 51 (45), 9124–9136. DOI: 10.1021/bi301039t From NLM Medline.

(52) Gumbart, J. C.; Roux, B.; Chipot, C. Efficient determination of protein-protein standard binding free energies from first principles. J Chem Theory Comput 2013, 9 (8). DOI: 10.1021/ct400273t From NLM PubMed-not-MEDLINE.

(53) Tidow, H.; Nissen, P. Structural diversity of calmodulin binding to its target sites. FEBS J 2013, 280 (21), 5551–5565. DOI: 10.1111/febs.12296 From NLM Medline.

(54) Samal, A. B.; Ghanam, R. H.; Fernandez, T. F.; Monroe, E. B.; Saad, J. S. NMR, biophysical, and biochemical studies reveal the minimal Calmodulin binding domain of the HIV-1 matrix protein. J Biol Chem 2011, 286 (38), 33533–33543. DOI: 10.1074/jbc.M111.273623 From NLM Medline.

(55) Osad’ko, I. S. Dependence of FRET efficiency on distance in single donor-acceptor pairs. J Chem Phys 2015, 142 (12), 125102. DOI: 10.1063/1.4915279 From NLM PubMed-not-MEDLINE.

(56) Park, S. H.; De Angelis, A. A.; Nevzorov, A. A.; Wu, C. H.; Opella, S. J. Three-dimensional structure of the transmembrane domain of Vpu from HIV-1 in aligned phospholipid bicelles. Biophys J 2006, 91 (8), 3032–3042. DOI: 10.1529/biophysj.106.087106 From NLM Medline.

(57) Volovik, M. V.; Denieva, Z. G.; Gifer, P. K.; Rakitina, M. A.; Batishchev, O. V. Membrane Activity and Viroporin Assembly for the SARS-CoV-2 E Protein Are Regulated by Cholesterol. Biomolecules 2024, 14 (9). DOI: 10.3390/biom14091061 From NLM Medline.

(58) Han, Z.; Harty, R. N. Influence of calcium/calmodulin on budding of Ebola VLPs: implications for the involvement of the Ras/Raf/MEK/ERK pathway. Virus Genes 2007, 35 (3), 511–520. DOI: 10.1007/s11262-007-0125-9 From NLM Medline.

(59) Banerjee, P.; Monje-Galvan, V.; Voth, G. A. Cooperative Membrane Binding of HIV-1 Matrix Proteins. J Phys Chem B 2024, 128 (11), 2595–2606. DOI: 10.1021/acs.jpcb.3c06222 From NLM Medline.

(60) Monette, A.; Niu, M.; Nijhoff Asser, M.; Gorelick, R. J.; Mouland, A. J. Scaffolding viral protein NC nucleates phase separation of the HIV-1 biomolecular condensate. Cell Rep 2022, 40 (8), 111251. DOI: 10.1016/j.celrep.2022.111251 From NLM Medline.

(61) Choi, S.; Lee, J. M.; Kim, K. K. Biomolecular condensates: molecular structure, biological functions, diseases, and therapeutic targets. Mol Biomed 2025, 6 (1), 99. DOI: 10.1186/s43556-025-00350-y From NLM Medline.

(62) Banani, S. F.; Lee, H. O.; Hyman, A. A.; Rosen, M. K. Biomolecular condensates: organizers of cellular biochemistry. Nat Rev Mol Cell Biol 2017, 18 (5), 285–298. DOI: 10.1038/nrm.2017.7 From NLM Medline.

(63) Corbi-Verge, C.; Kim, P. M. Motif mediated protein-protein interactions as drug targets. Cell Commun Signal 2016, 14, 8. DOI: 10.1186/s12964-016-0131-4 From NLM Medline.

(64) Caporale, A.; Adorinni, S.; Lamba, D.; Saviano, M. Peptide-Protein Interactions: From Drug Design to Supramolecular Biomaterials. Molecules 2021, 26 (5). DOI: 10.3390/molecules26051219 From NLM Medline.

(65) Chadda, R.; Lee, T.; Mahoney-Kruszka, R.; Kelley, E. G.; Bernhardt, N.; Sandal, P.; Robertson, J. L. A thermodynamic analysis of CLC transporter dimerization in lipid bilayers. Proceedings of the National Academy of Sciences 2023, 120 (41), e2305100120.

(66) Jiang, L.; Xiong, Z.; Song, Y.; Lu, Y.; Chen, Y.; Schultz, J. S.; Li, J.; Liao, J. Protein– protein affinity determination by quantitative FRET quenching. Scientific reports 2019, 9 (1), 2050.

(67) Shrestha, D.; Jenei, A.; Nagy, P.; Vereb, G.; Szöllősi, J. Understanding FRET as a research tool for cellular studies. International journal of molecular sciences 2015, 16 (4), 6718–6756.

(68) Yi, S.; Kim, E.; Yang, S.; Kim, G.; Bae, D. W.; Son, S. Y.; Jeong, B. G.; Ji, J. S.; Lee, H. H.; Hahn, J. S. Direct quantification of protein–protein interactions in living bacterial cells. Advanced Science 2025, 12 (19), 2414777.

