## Supplementary material for "Quantitative and amino acid sequence analysis of soluble HIV-1 Vpu and calmodulin interactions": Manuscript

[#]Equal contribution


**A SUMO-FL Vpu**

MGSDSEVNQEAKPEVKPEVKPETHINLKVSDGSSEIFFKIKKTTPLRRLMEAFAKRQGKE
MDSLRFLYDGIRIQADQTPEDLDMEDNDIIEAHREQIGGSPSGNQGSGLVPRGS**SGQGS**M
QPIPIVAIVALVVAIIIAIV**VW**SIVIIEYRK**I**LRCRKIDRLIDRLIERAEDSGNESEGEI
SALVEMGVEMGHHAPWDVDDLSGHHHHHHHHHH

**B SUMO-Vpu C-terminal region (residues 28-81)**

MGHHHHHHHHGSNGMSDSEVNQEAKPEVKPEVKPETHINLKVSDGSSEIFFKIKKTTPLR
RLMEAFAKRQGKEMDSLRFLYDGIRIQADQTPEDLDMEDNDIIEAHREQIGGSPSGNQGS
GLVPRGS**SGQGSG**MEYRKILRCRKIDRLIDRLIERAEDSGNESEGEISALVEMGVEMGHH
APWDVDDL

**Figure S1.** Amino acid sequence of the SUMO-tagged Vpu constructs—FL WT and V22A/W23Y mutant Vpu (A), as well as Vpu's C-terminal region (B) were produced as chimera constructs with a SUMO tag fused to their N-termini. Poly-histidine (His8 or His10) tag at either N- or C-terminus was used for affinity purification. The thrombin-cutting site is highlighted in red. The L42C used for fluorescent labeling with Cy3 dye is highlighted in cyan. The residues V22,W23 and I33 substituted by A, Y, and N, respectively, in the Vpu-M construct are bold blue. The linker between the Vpu variant and the Thrombin-recognition site are underlined and in bold. These linker residues were present in the FL WT Vpu and FL Vpu-M after the SUMO tag was removed via Thrombin digestion.

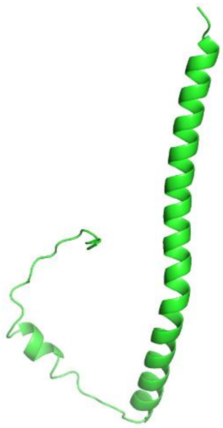

**Figure S2.** AlphaFold model of FL Vpu with the V22A/W23Y/I33N mutations. The structure is very close to those predicted for FL WT Vpu[1]. AlphaFold in Chimera 1.11.1[2] was used. For both WT FL Vpu and FL Vpu-M, the predicted structures deviate from the NMR-determined structure of FL Vpu in lipid[3]. This NMR structure is shown in the main text in Figure 1A.

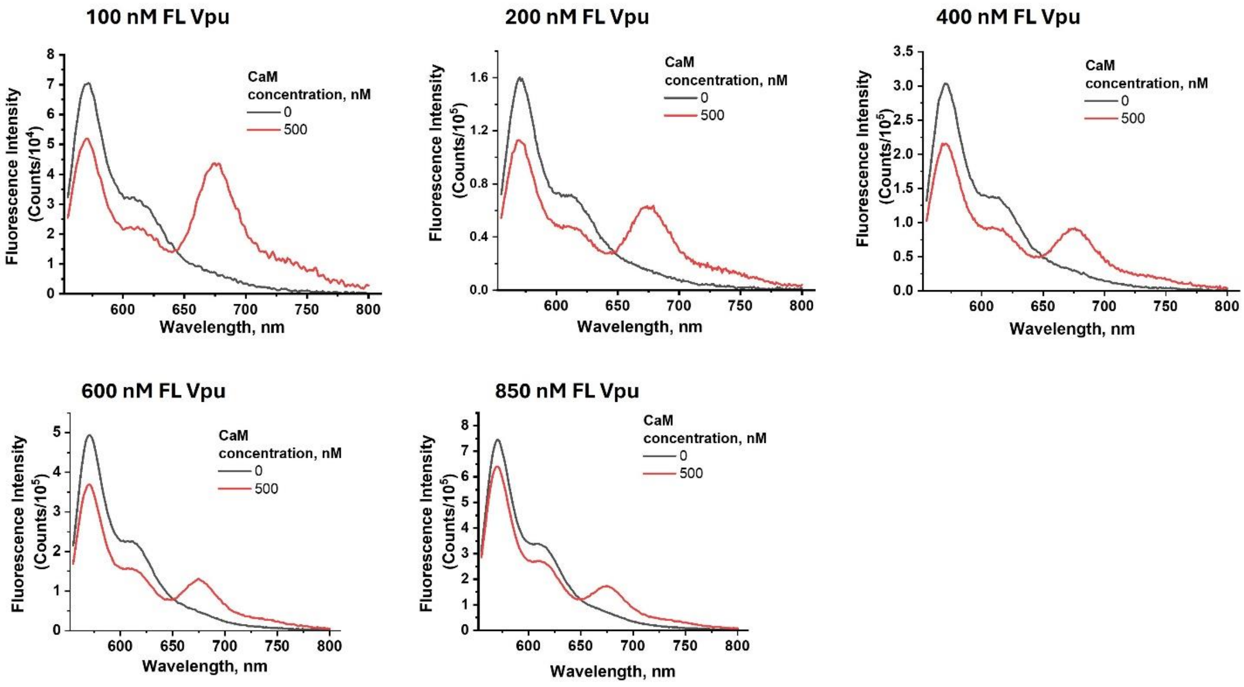


**Figure S3.** The raw eFRET data between Cy5-labeled $Ca^{2+}$-CaM at 500 nM constant concentration and increasing concentration of Cy3-labeled FL WT Vpu in a range from 100 nM to 850 nM. Each FRET data shows the signal at 0 nM (black) and 500 nM Cy5-labeled $Ca^{2+}$-CaM (red).

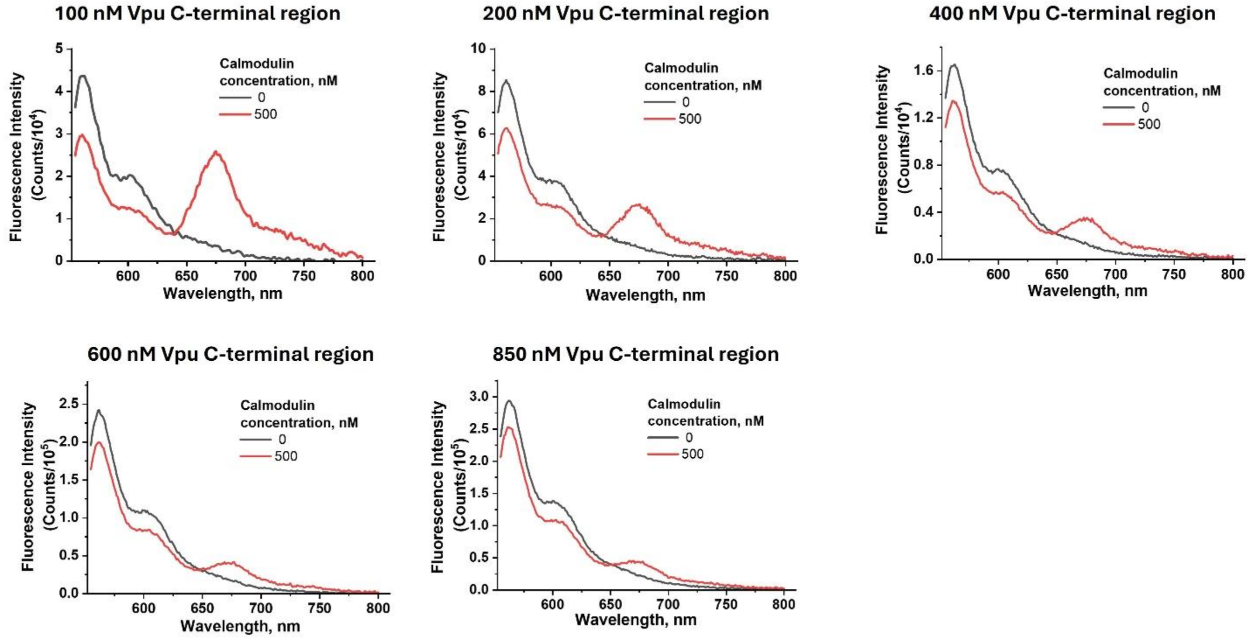


**Figure S4**. The raw eFRET data between constant concentration of Cy5-labeled $Ca^{2+}$-CaM (500 nM) and increasing concentration of Cy3-labeled CT Vpu in a range from 100 nM to 850 nM. Each FRET data shows the signal at 0 nM (black) and 500 nM Cy5-labeled $Ca^{2+}$-CaM (red).

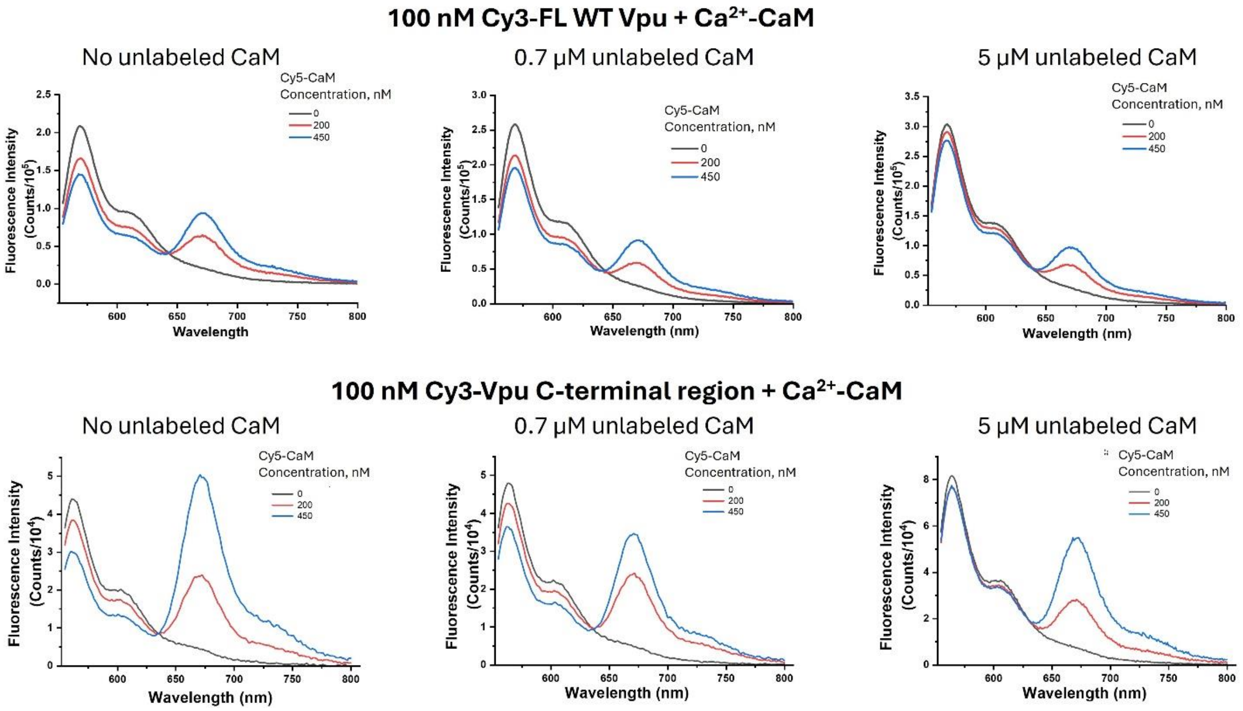


**Figure S5.** Raw eFRET data for the Cy3-Vpu-Cy5-$Ca^{2+}$-CaM without and with competitive binding of non-labeled (unlabeled) $Ca^{2+}$-CaM. The data for Cy3-FL WT Vpu and Cy3-Vpu C-terminal region are shown in (A) and (B), respectively. The more intensive fluorescence emission of Cy3-$Ca^{2+}$-CaM when bound to Cy3-Vpu C-terminal compared to Cy3-FL WT Vpu is most likely a result of conformational arrangements and proximity of the Cy3 donor and Cy5 acceptor.

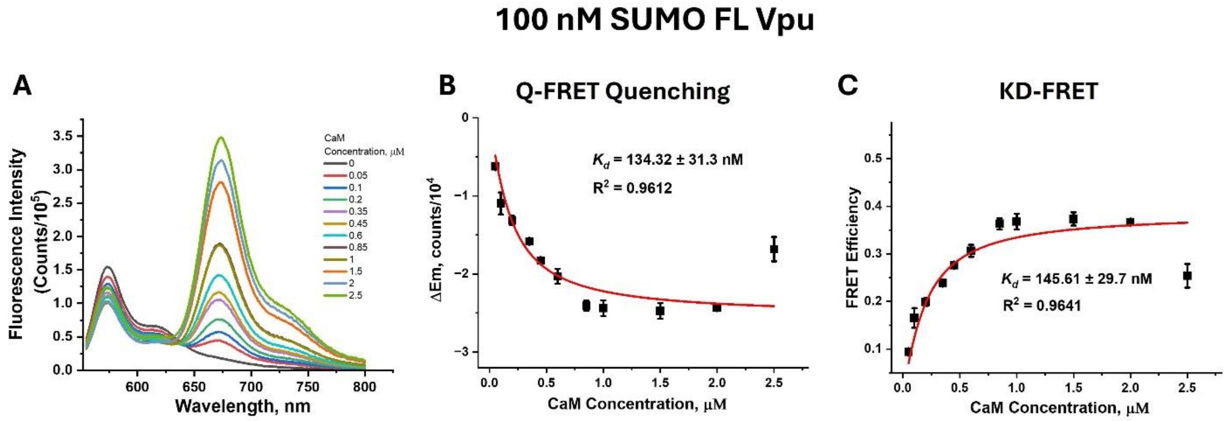


**Figure S6.** eFRET data and analyses for the interaction of 100 nM Cy3-SUMO-FL WT Vpu with Cy5-$Ca^{2+}$-CaM at increasing concentration in the range from 0 to 2.5 μM. The original eFRET data are shown in (A)—The Cy3 emission decreases while Cy5 emission increases upon Cy3-Vpu-$Ca^{2+}$-CaM complex formation. The background-corrected eFRET data (in A) were used to estimate $K_d$. The results from fitting the Cy3 donor fluorescence decay (method of Quantitative FRET Quenching) and fitting the FRET efficiencies ($E_{FRET}$) (KD-FRET) to obtain $K_d$-s are shown

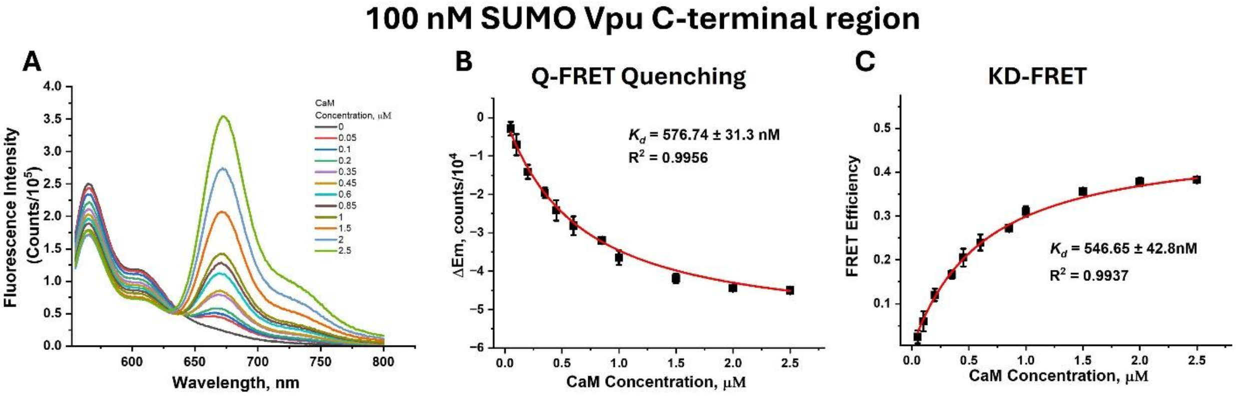


**Figure S7.** eFRET data and analyses for the interaction of 100 nM Cy3-SUMO-Vpu C-terminal with Cy5-$Ca^{2+}$-CaM at increasing concentration in the range from 0 to 2.5 μM. The original eFRET data are shown in (A)—The Cy3 emission decreases while Cy5 emission increases upon Cy3-Vpu-$Ca^{2+}$-CaM complex formation. The background-corrected eFRET data (in A) were used to estimate $K_d$. The results from fitting the Cy3 donor fluorescence decay (method of Quantitative FRET Quenching) and fitting the FRET efficiencies ($E_{FRET}$) (KD-FRET) to obtain $K_d$-s are shown in (B) and (C), respectively. The estimated $K_d$-s and quality of data fits are shown in (B) and (C) inset. All data are average of triplicate samples.

### Supporting References


1. Ishola, O; Islam, MM; Hadadianpour, E; Borbat, PP; Ogunbowale, A; Amador, JCR; Georgieva, ER. The soluble state of the HIV-1 Vpu protein forms a complex with Ca2+-calmodulin. *Protein Science*. (**2026**), *35*, e70487. DOI: doi: https://doi.org/10.1002/pro.70487Digital

2. Mirdita, M; Schutze, K; Moriwaki, Y; Heo, L; Ovchinnikov, S; Steinegger, M. ColabFold: making protein folding accessible to all. *Nat Methods*. (**2022**), *19* (6), 679-682. DOI: 10.1038/s41592-022-01488-1

3. Zhang, H; Lin, EC; Das, BB; Tian, Y; Opella, SJ. Structural determination of virus protein U from HIV-1 by NMR in membrane environments. *Biochim Biophys Acta*. (**2015**), *1848* (11 Pt A), 3007-3018. DOI: 10.1016/j.bbamem.2015.09.008
